# Two forms of phosphomannomutase in gammaproteobacteria: The overlooked membrane-bound form of AlgC is required for twitching motility of *Lysobacter enzymogenes*

**DOI:** 10.1101/589796

**Authors:** Guoliang Qian, Shifang Fei, Michael Y. Galperin

## Abstract

*Lysobacter enzymogenes*, a member of *Xanthomonadaceae*, is a promising tool to control crop-destroying fungal pathogens. One of its key antifungal virulence factors is the type IV pili that are required for twitching motility. Transposon mutagenesis of *L*. *enzymogenes* revealed that production of type IV pili required the presence of the *Le2152* gene, which encodes an AlgC-type phosphomannomutase/phosphoglucomutase (PMM). However, in addition to the cytoplasmic PMM domain, the Le2152 gene product contains a ca. 200-aa N-terminal periplasmic domain that is anchored in the membrane by two transmembrane segments and belongs to the dCache superfamily of periplasmic sensor domains. Sequence analysis identified similar membrane-anchored PMMs, encoded in conserved *coaBC*-*dut*-*algC* gene clusters, in a variety of gammaproteobacteria, either as the sole PMM gene in the entire genome or in addition to the gene encoding the stand-alone enzymatic domain. Previously overlooked N-terminal periplasmic sensor domains were detected in the well-characterized PMMs of *Pseudomonas aeruginosa* and *Xanthomonas campestris*, albeit not in the enzymes from *Pseudomonas fluorescens, Pseudomonas putida* or *Azotobacter vinelandii*. It appears that after the initial cloning of the enzymatically active soluble part of *P*. *aeruginosa* AlgC in 1991, all subsequent studies utilized N-terminally truncated open reading frames. The N-terminal dCache sensor domain of AlgC is predicted to modulate the PMM activity of the cytoplasmic domain in response to as yet unidentified environmental signal(s). AlgC-like membrane-bound PMMs appear to comprise yet another environmental signaling system that regulates production of type IV pili and potentially other systems in certain gammaproteobacteria.

## INTRODUCTION

*Lysobacter enzymogenes*, a member of the gammaproteobacterial family *Xanthomonadaceae*, is a promising organism for biocontrol of fungal plant pathogens (Zhang and Yuen, 1999; Qian *et al*., 2009). It infects filamentous fungal pathogens, such as *Bipolaris sorokiniana*, the causative agent of common root rot and spot blotch of barley and wheat seeds, and suppresses their growth through secretion of a heat-stable antifungal factor and production of extracellular chitinase, lysobactin, and other compounds (Zhang and Yuen, 2000; Yu *et al*., 2007; Li *et al*., 2008; de Bruijn *et al*., 2015). Colonization by *L*. *enzymogenes* depends on formation of type IV pili (T4P) and T4P-mediated twitching motility (Patel *et al*., 2010; 2011). Twitching motility mediated by T4P is considered an important taxonomical feature for the genus *Lysobacter* (Christensen and Cook, 1978).

Using *L*. *enzymogenes* strain OH11, originally isolated from the rhizosphere of green pepper (Qian *et al*., 2009), as a working model, we have previously identified transcriptional regulators Clp and PilR as key regulators of T4P-mediated twitching motility (Wang *et al*., 2014; Chen *et al*., 2017; Chen *et al*., 2018). Aiming to identify new regulators modulating twitching motility, we have conducted transposon mutagenesis of *L*. *enzymogenes* strain OH11 and analyzed the open reading frames (ORFs) whose disruption abolished twitching motility. Here we describe one of such ORFs, Le2152, which encodes a membrane-bound phosphomannomutase with an N-terminal periplasmic sensor domain, and analyze the distribution of such two-domain phosphomannomutases.

## RESULTS

### Le2152 protein is required for type IV pili-mediated twitching motility

We have used a wild-type environmental isolate of *L*. *enzymogenes*, strain OH11 (Qian *et al*., 2009), to conduct transposon mutagenesis with Tn5 and generate a mutant library. After screening more than 300 mutant strains, we identified a mutant that completely lacked twitching motility. The disrupted gene in this mutant was identified as *Le2152* (locus tag D9T17_01580, GenBank accession number ROU09072.2), which encodes a 772-aa protein that was predicted to function as a phosphomannomutase. To validate the role of *Le2125* in twitching motility, we generated an in-frame deletion mutant via homologous double cross-over recombination, as described previously (Qian *et al*., 2012), see Supporting Information Tables S1 and S2 for experimental details. As shown in Figure 1, this in-frame deletion mutant, Δ*Le2152*, was also deficient in twitching motility, as no mobile cells were observed at the margin of its colony, whereas wild-type OH11 had numerous mobile cells at the margin of its colonies. Introduction of a plasmid-borne *Le2152* gene under its native promoter fully restored twitching motility of Δ*Le2152*, while the Δ*Le2152* strain carrying an empty vector was still deficient in this function. In contrast to the Δ*Le2152* strain, a mutant with a deletion in *Le4861* gene that codes for a closely related enzyme phosphoglucosamine mutase (GlmM, locus tag D9T17_13225, GenBank accession ROU06416.1) exhibited no motility defect (Supporting Information Figure S1). These results show that the product of *Le2152* is required for twitching motility in *L*. *enzymogenes*.

**Figure 1.**
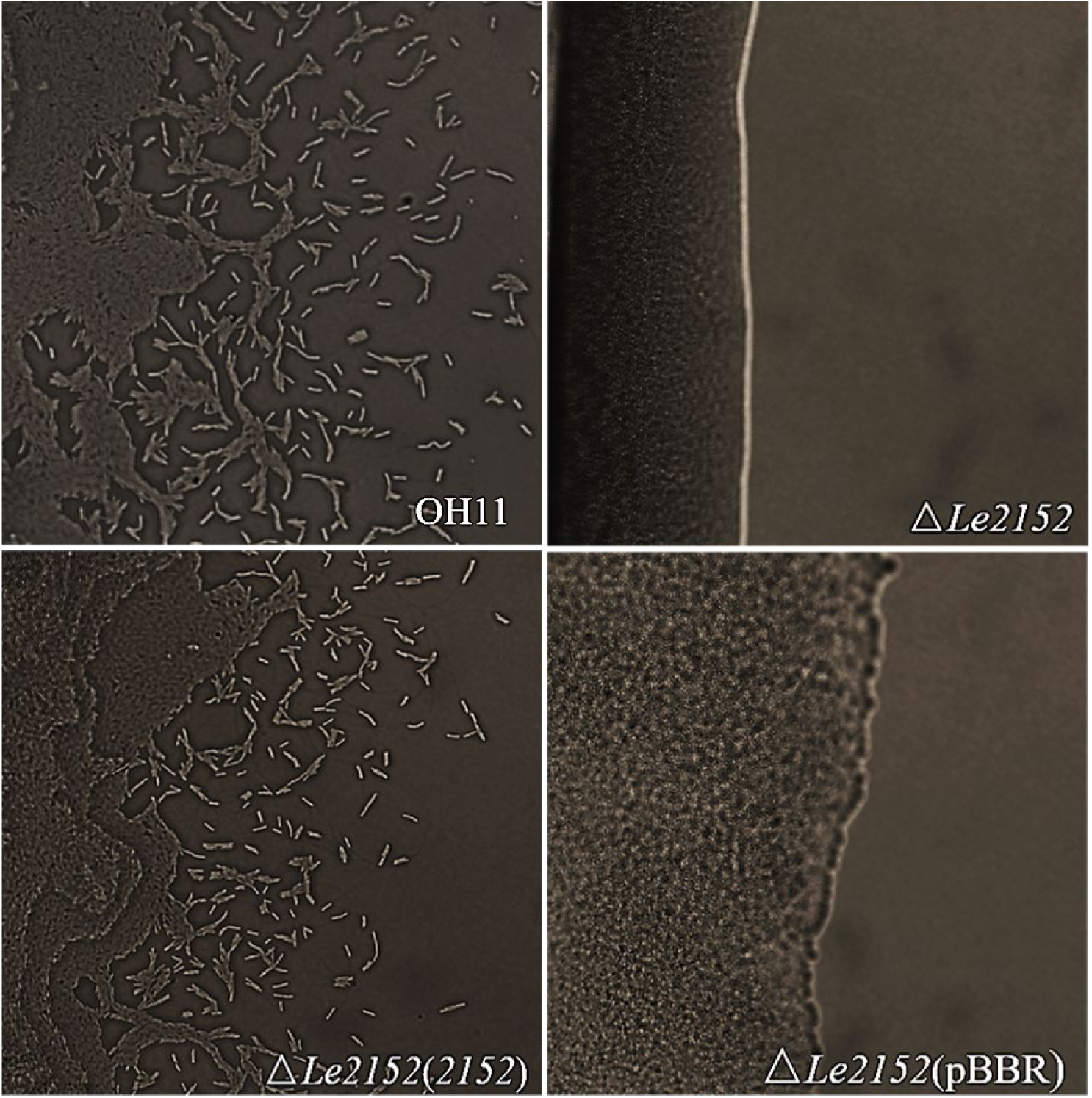
Le2152 is required for twitching motility in Lysobacter enzymogenes OH11. Loss of *Le2152* resulted in the absence of motile cells on the margins of the mutant colonies. **A**. OH11, wild-type strain of *L*. *enzymogenes*. **B**. Δ*Le2152*, the *Le2152* in-frame deletion mutant of OH11. **C**. Δ*Le2152*(*2152*), Δ*Le2152* strain complemented with plasmid-borne *Le2152* under its native promoter. **D**. Δ*Le2152*(pBBR), Δ*Le2152* strain haboring an empty vector.

### Domain architecture and phylogenetic distribution of Le2152 homologs

Sequence analysis of the *Le2152* gene product revealed that it contains a typical 450-aa cytoplasmic phosphomannomutase (PMM) domain. However, this domain is preceded by a 320-aa N-terminal fragment that consists of 200-aa predicted periplasmic domain, which is anchored in the membrane by two transmembrane segments and followed by a 55-aa Pro-rich flexible linker (Figure 2).

**Figure 2.**
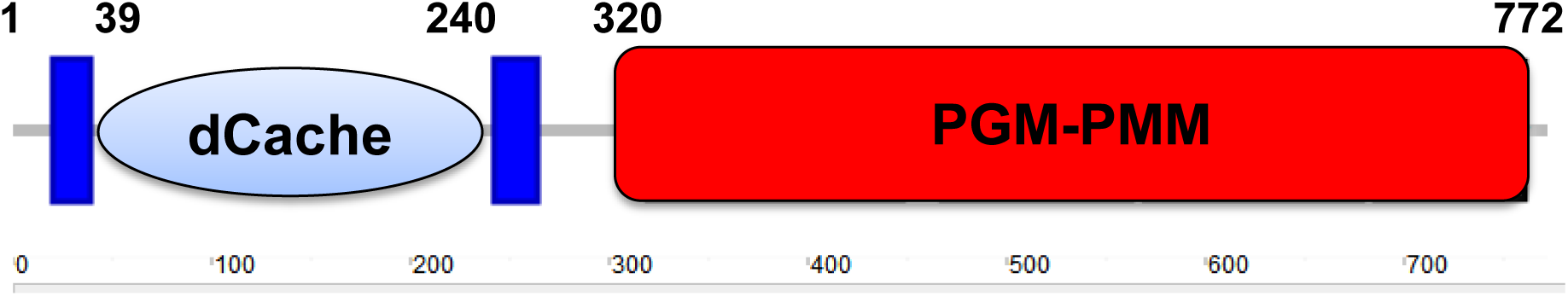
Domain organization of the Lysobacter enzymogenes Le2152 protein. Blue boxes indicate predicted transmembrane regions; PGM-PMM indicates the cytoplasmic phosphomannomutase domain, a combination of Pfam (El-Gebali *et al*., 2019) domains PGM_PMM_I (PF02878), PGM_PMM_II (PF02879), PGM_PMM_III (PF02880), and PGM_PMM_IV (PF00408). The transmembrane orientation and domain architecture of Le2152 were predicted using the TMHMM and SMART tools (Krogh *et* al., 2001; Letunic and Bork, 2018). The predicted periplasmic domain was identified as dCache using HHpred (Zimmermann *et al*., 2018), see Supporting Information Figure S2 for details.

To check if the full-length Le2152 protein – or just its cytoplasmic enzymatic domain – is expressed *in vivo*, we expressed in *L*. *enzymogenes* the Le2152 protein fused with a C-terminal FLAG tag from the pBBR1-MCS5 plasmid under its native promoter and performed a Western blot, probing it with anti-FLAG antibodies. The expressed Le2152 protein showed molecular weight of ca. 100 KDa, which is above the calculated molecular weight of 82 kDa of the full-length Le2152 (Supporting Information Figure S2A). The reason for this discrepancy is still unclear, but this observation shows that the full-length Le2152 is indeed expressed *in vivo* and is the form that is essential for the *L*. *enzymogenes* twitching motility.

Sensitive sequence similarity searches with HHPred (Zimmermann *et al*., 2018) allowed assigning the N-terminal periplasmic domain of Le2152 to the Double Cache (dCache) domain superfamily (Upadhyay *et al*., 2016), previously referred to as PDC (PhoQ, DcuS and CitA) fold domains (Cheung *et al*., 2008; Zhang and Hendrickson, 2010; Pineda-Molina *et al*., 2012). Within the dCache superfamily, the periplasmic domain of Le2152 showed the highest similarity to the Pfam domain *dCache_3* (PF14827, (El-Gebali *et al*., 2019)). Among domains of known 3D structure, it was most similar to the *dCache_1* (PF02743) sensor domain of the *Bacillus subtilis* sporulation kinase KinD (Protein DataBank entry 4JGP) and C4-dicarboxylate-binding sensor domains of the histidine kinase DctB from *Vibrio cholerae* and *Sinorhizobium meliloti* (PDB: 3BY9 and 3E4P) (Cheung and Hendrickson, 2008; Zhou *et al*., 2008), which are described in Pfam as the C*ache_3-Cache_2* fusion domain (PF17201). Thus, the periplasmic domain of Le2152 falls somewhere in-between *dCache_1, dCache_3*, and C*ache_3-Cache_2* domains and likely represents a new family of dCache domains, which could be a reason why it has escaped recognition for so long. The sequence logo and a representative alignment of the dCache domain of Le2152 are shown in Supporting Information Figures S2. This figure also shows a sequence logo of the flexible linker domain and an alignment of the cytoplasmic enzymatic domain of Le2152.

Sequence similarity searches using Le2152 protein as the query revealed the presence of similar two-domain membrane-anchored PMMs in a variety of gammaproteobacteria. Out of the currently recognized 20 orders of gammaproteobacteria, such proteins were found to be encoded in representatives of at least eight: *Alteromonadales, Cellvibrionales, Chromatiales, Methylococcales, Oceanospirillales, Pseudomonadales, Thiotrichales*, and *Xanthomonadales* (Table 1). Many of these bacteria, including the well-studied model organisms *Alcanivorax borkumensis, Stenotrophomonas maltophilia* and *Xanthomonas citri*, carry two PMM genes: one (the “long” version) that codes for the two-domain membrane-anchored PMM and the other (the “short” version) that codes for the stand-alone enzymatic domain. In other organisms, such as *Lysobacter spp*., the membrane-anchored PMM is the only one encoded in the genome, although all these organisms also carry the *glmM* genes that code for the closely related soluble phosphoglucosamine mutase (Supporting Information Table S3), which has been reported to have certain PMM activity (Tavares *et al*., 2000).

**Table 1.**
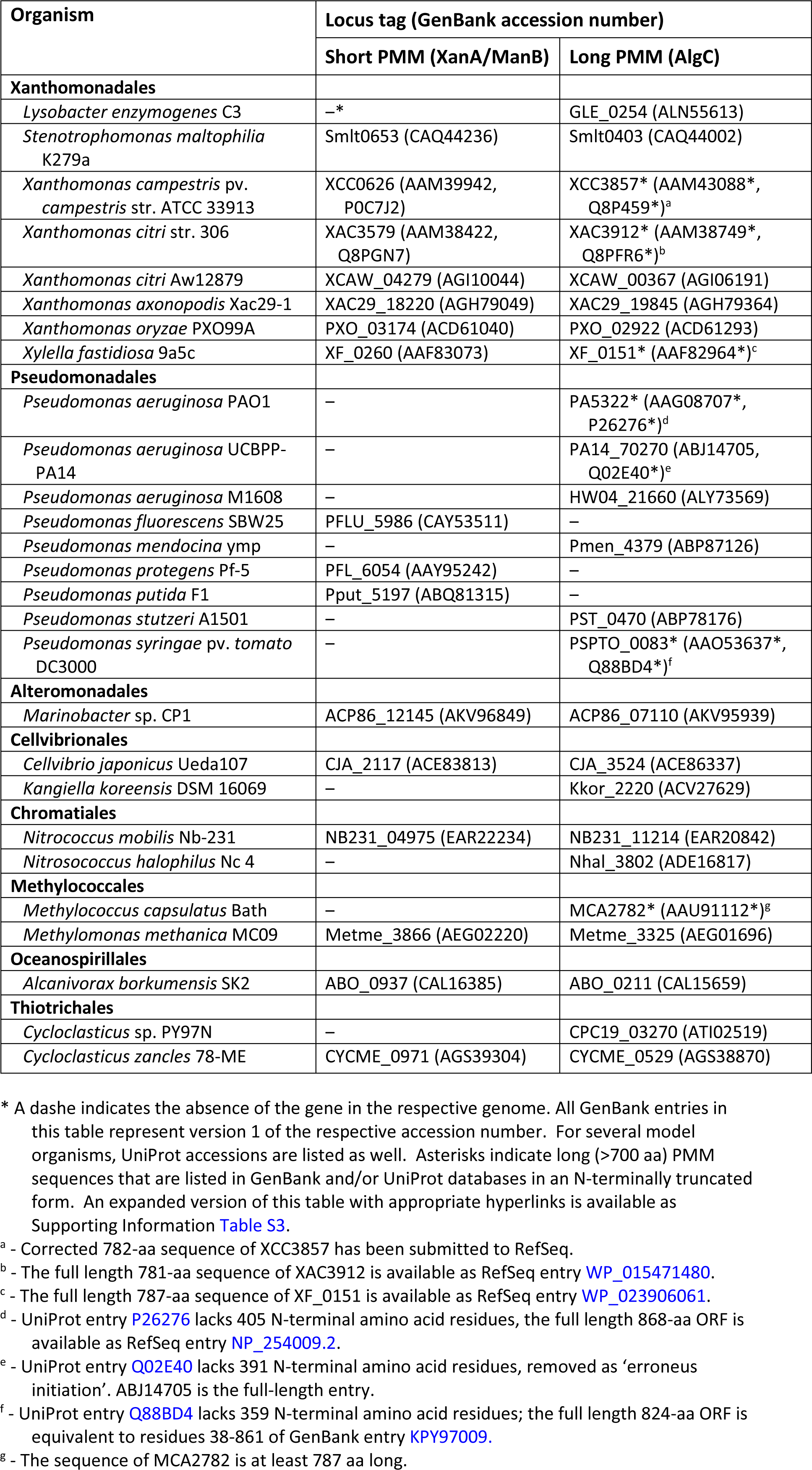
Two types of phosphomannomutase in gammaproteobacteria

| Organism | Locus tag (GenBank accession number) |  |
| --- | --- | --- |
|  | Short PMM (XanA/ManB) | Long PMM (AlgC) |
| **Xanthomonadales** |  |  |
| *Lysobacter enzymogenes* C3 | –* | GLE_0254 (ALN55613) |
| *Stenotrophomonas maltophilia* K279a | Smlt0653 (CAQ44236) | Smlt0403 (CAQ44002) |
| *Xanthomonas campestris* pv. *campestris* str. ATCC 33913 | XCC0626 (AAM39942, P0C7J2) | XCC3857* (AAM43088*, Q8P459*)<sup>a</sup> |
| *Xanthomonas citri* str. 306 | XAC3579 (AAM38422, Q8PGN7) | XAC3912* (AAM38749*, Q8PFR6*)<sup>b</sup> |
| *Xanthomonas citri* Aw12879 | XCAW_04279 (AGI10044) | XCAW_00367 (AGI06191) |
| *Xanthomonas axonopodis* Xac29-1 | XAC29_18220 (AGH79049) | XAC29_19845 (AGH79364) |
| *Xanthomonas oryzae* PXO99A | PXO_03174 (ACD61040) | PXO_02922 (ACD61293) |
| *Xylella fastidiosa* 9a5c | XF_0260 (AAF83073) | XF_0151* (AAF82964*)<sup>c</sup> |
| **Pseudomonadales** |  |  |
| *Pseudomonas aeruginosa* PAO1 | – | PA5322* (AAG08707*, P26276*)<sup>d</sup> |
| *Pseudomonas aeruginosa* UCBPP-PA14 | – | PA14_70270 (ABJ14705, Q02E40*)<sup>e</sup> |
| *Pseudomonas aeruginosa* M1608 | – | HW04_21660 (ALY73569) |
| *Pseudomonas fluorescens* SBW25 | PFLU_5986 (CAY53511) | – |
| *Pseudomonas mendocina* ymp | – | Pmen_4379 (ABP87126) |
| *Pseudomonas protegens* Pf-5 | PFL_6054 (AAY95242) | – |
| *Pseudomonas putida* F1 | Pput_5197 (ABQ81315) | – |
| *Pseudomonas stutzeri* A1501 | – | PST_0470 (ABP78176) |
| *Pseudomonas syringae* pv. *tomato* DC3000 | – | PSPTO_0083* (AAO53637*, Q88BD4*)<sup>f</sup> |
| **Alteromonadales** |  |  |
| *Marinobacter* sp. CP1 | ACP86_12145 (AKV96849) | ACP86_07110 (AKV95939) |
| **Cellvibrionales** |  |  |
| *Cellvibrio japonicus* Ueda107 | CJA_2117 (ACE83813) | CJA_3524 (ACE86337) |
| *Kangiella koreensis* DSM 16069 | – | Kkor_2220 (ACV27629) |
| **Chromatiales** |  |  |
| *Nitrococcus mobilis* Nb-231 | NB231_04975 (EAR22234) | NB231_11214 (EAR20842) |
| *Nitrosococcus halophilus* Nc 4 | – | Nhal_3802 (ADE16817) |
| **Methylococcales** |  |  |
| *Methylococcus capsulatus* Bath | – | MCA2782* (AAU91112*)<sup>g</sup> |
| *Methylomonas methanica* MC09 | Metme_3866 (AEG02220) | Metme_3325 (AEG01696) |
| **Oceanospirillales** |  |  |
| *Alcanivorax borkumensis* SK2 | ABO_0937 (CAL16385) | ABO_0211 (CAL15659) |
| **Thiotrichales** |  |  |
| *Cycloclasticus* sp. PY97N | – | CPC19_03270 (ATI02519) |
| *Cycloclasticus zancles* 78-ME | CYCME_0971 (AGS39304) | CYCME_0529 (AGS38870) |

\* A dashe indicates the absence of the gene in the respective genome. All GenBank entries in this table represent version 1 of the respective accession number. For several model organisms, UniProt accessions are listed as well. Asterisks indicate long (>700 aa) PMM sequences that are listed in GenBank and/or UniProt databases in an N-terminally truncated form. An expanded version of this table with appropriate hyperlinks is available as Supporting Information Table S3. <sup>a</sup> - Corrected 782-aa sequence of XCC3857 has been submitted to RefSeq. <sup>b</sup> - The full length 781-aa sequence of XAC3912 is available as RefSeq entry WP\_015471480. <sup>c</sup> - The full length 787-aa sequence of XF\_0151 is available as RefSeq entry WP\_023906061. <sup>d</sup> - UniProt entry P26276 lacks 405 N-terminal amino acid residues, the full length 868-aa ORF is available as RefSeq entry NP\_254009.2. <sup>e</sup> - UniProt entry Q02E40 lacks 391 N-terminal amino acid residues, removed as 'erroneous initiation'. ABJ14705 is the full-length entry. <sup>f</sup> - UniProt entry Q88BD4 lacks 359 N-terminal amino acid residues; the full length 824-aa ORF is equivalent to residues 38-861 of GenBank entry KPY97009. <sup>g</sup> - The sequence of MCA2782 is at least 787 aa long.

The “short” PMM genes have variable genomic neighborhoods that often include the *xanB* (or *cpsB*) gene that codes for the bifunctional mannose-1-phosphate guanylyltransferase/phosphomannose isomerase. By contrast, the “long” PMM genes reside in conserved gene clusters that include the *coaBC* gene(s) encoding bifunctional phosphopantothenoylcysteine decarboxylase and phosphopantothenate–cysteine ligase; the *dut* gene, which encodes the house-cleaning dUTPase, and, in many organisms, the N-acetylglutamate kinase gene *argB* (Figure 3, see also Supporting Information Figure S3).

**Figure 3.**
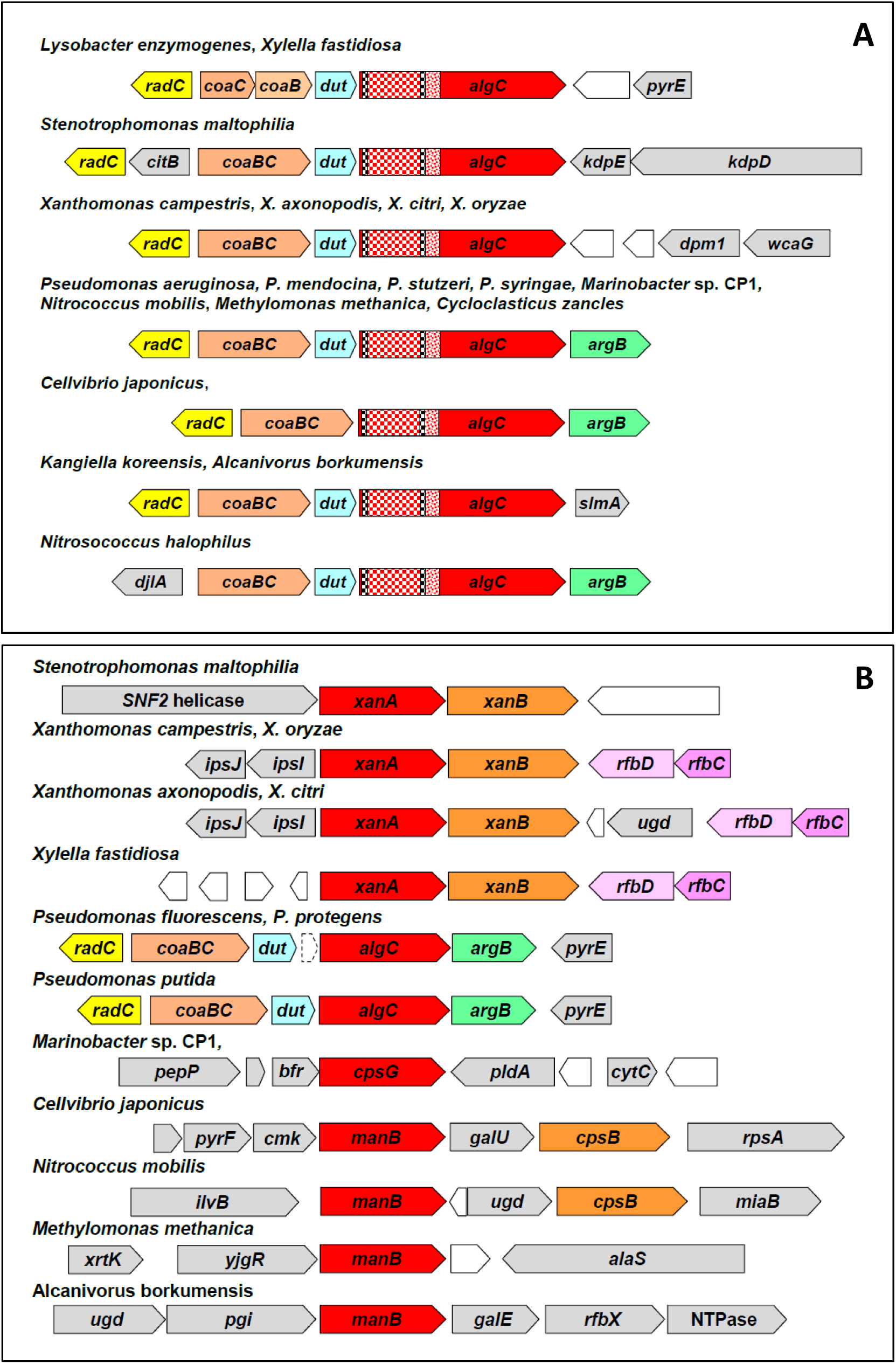
Genomic neighborhoods of phosphomannomutase genes in selected gammaproteobacteria. Phosphomannomutase (PMM) genes are shown in red, their genomic locus tags and GenBank accession numbers are listed in Table 1. Members of the conserved neighborhoods are indicated with bright colors with same colors for all homologs; variable genes are in grey, uncharacterized genes are in white. Gene names are from the COG database (Galperin *et al*., 2015), the shapes are drawn approximately to size. **A**. Gene clusters of the *algC* (“long” PMM) genes. The coloring of *algC* reflects domain organization of its product: the periplasmic dCache domain is shown as red checkered box, two transmembrane segments as black checkered boxes, the flexible linker as red dotted box and the enzymatic domain is in red. **B**. Gene clusters of the “short” PMM genes. See text for details.

All long PMMs, retrieved by iterative sequence similarity searches with PSI-BLAST and jackHMMer, were predicted to have essentially the same domain architecture as Le2152 (Figure 2), which included a dCache-type periplasmic sensor domain anchored by two transmembrane helices and followed by a flexible linker and the cytoplasmically located enzymatic domain. These (predicted) periplasmic domains displayed only a limited sequence similarity with few conserved residues (Supporting Information Figure S2B), consistent with a ligand-binding, rather than enzymatic, function.

Remarkably, in the original genome annotations of *Xylella fastidiosa, Xanthomonas campestris*, and *X*. *citri*, translations of the membrane-bound PMMs were artificially truncated by removing the N-terminal fragments and leaving each of these organisms with two ORFs encoding only the XanA/ManB-like enzymatic domain (Table 1). In subsequent annotations, some of the full-length ORFs have been restored but N-terminally truncated PMMs are still listed in certain GenBank and UniProt entries (Supporting Information Table S3 and Figure S3).

Just like *Lysobacter* spp., all checked members of the *Pseudomonas* genus carried only a single PMM gene. However, while some *Pseudomonas* spp. encoded the membrane-anchored two-domain version of the enzyme, others encoded only the XanA-like cytoplasmic version (Table 1, Figure 3). Surprisingly, the N-terminal periplasmic dCache domain was also detected in the ORFs of the well-characterized phosphomannomutase AlgC of *P*. *aeruginosa* strains PAO1 and PA14 (Table 1). While the enzymatically active C-terminal part of this protein had been cloned back in 1991 (Zielinski *et al*., 1991), the full-length 868-aa ORF was only translated from the complete genome sequence. Still, the shorter 463-aa ORF was routinely assumed to be the correct one. With almost 2,000 genome sequences of various strains of *P*. *aeruginosa* available in the public databases as of 01.01.2019, GenBank contained 1,785 sequences of the two-domain AlgC (ranging in length from 863 to 870 aa) and 26 sequences of the enzymatically active shorter version of this protein (from 463 to 470 aa). In *Pseudomonas syringae*, about half of the annotated PMM sequences from various strains were of the “short” variety (usually 465 aa long) but these genes were always preceded by an untranslated region of ∼1.2 kb. By contrast, all PMM genes from *Pseudomonas fluorescens, Pseudomonas protegens*, and *Pseudomonas putida* were of the “short” variety and had no significant gaps in front of them (Figure 3, see also Supporting Information Figure S3). However, these short PMM genes of *Pseudomonas* spp. were still located in the same conserved *coaBC*-*dut*-*algC*-*argB* gene clusters as the long genes. Further, while most C-terminal enzymatic domains of long and short PMMs clustered separately from each other, forming two well-resolved clades, products of short PMM genes of *P*. *fluorescens, P*. *protegens* and *P*. *putida* clustered with the long PMMs from other pseudomonads, rather than with short PMMs from other organisms (Supporting Information Figure S4). These observations suggest that (i) the short PMMs of *Pseudomonas* spp. evolved through the loss of their N-terminal fragments and (ii) this loss was a relatively recent event.

### Expression of the two-domain phosphomannomutase

Given that only the C-terminal parts of long PMMs have the enzymatic activity, the question arises if the full-size ORFs are getting expressed. As mentioned above, the full-length Le2152 protein expressed from its native promoter exhibited molecular weight of ca. 100 kDa (Supporting Information Figure S2A). In addition, examination of the publicly available transcriptomic data for several “long” PMM-encoding bacteria, available in the NCBI’s Sequence Read Archive (SRA), revealed expression of the RNAs corresponding to the N-terminal dCache domains of their PMMs (Supporting Information Figure S5). In addition to *L*. *enzymogenes*, transcribed RNAs from the 5’ regions of the respective ORFs have been found in the SRA entries for various *Xanthomonas* spp., including *X*. *campestris* strains 8004 and B100 (Bonomi *et al*., 2016; Wang *et al*., 2017; Alkhateeb *et al*., 2018), *X*. *citri* (Jalan *et al*., 2013), and *X*. *oryzae* (Kim *et al*., 2016), as well as *S*. *maltophilia* strains K279a and FLR (Abda *et al*., 2015; Gallagher *et al*., 2019) and *Xylella fastidiosa* strain ‘Temecula’ (Parker *et al*., 2016), see Supporting Information Table S4. A similar picture, showing expression of the full-length AlgC-type PMM, was observed in *P*. *aeruginosa* strains PAO1 and PA14 (Supporting Information Figure S5), although in the former several SRA profiles showed expression of only the cytoplasmic fragment. Expression of “long” PMM genes was also detected in other Pseudomonas spp., as well as in representatives of other gammaproteobacterial orders, such as *Cellvibrio japonicus* (Blake *et al*., 2018) and *Methylobacter tundripaludum* (Krause *et al*., 2017). Finally, unpublished RNA-Seq data from the DOE Joint Genome Institute revealed robust expression of the full-length PMM gene of *Methylococcus capsulatus*, which is currently listed in GenBank in truncated form (Supporting Information Table S4). These data clearly show the presence of transcripts for the N-terminal dCache domain of “long” PMMs in a variety of gammaproteobacteria. At the same time, these RNA-Seq profiles often showed an even larger number of transcripts for the C-terminal enzymatic parts of these PMMs. Taken together, these data indicate that the full-size PMM genes are actively transcribed and are subject to a complex regulation with the possibility of additional transcription starts.

## DISCUSSION

A widespread group of experimentally characterized bacterial genes, referred to as *algC* in *P*. *aeruginosa, exoC* in *Azospirillum brasilense, manB* (*cpsG*) *and rfbK* in *Escherichia coli* and *Salmonella, noeK* in *Sinorhizobium fredii, pgmG* in *Sphingomonas sanxanigenens, pmmA* in *Prochlorothrix hollandica, rfbB* in *Vibrio cholerae, spgM* in *S*. *maltophilia* and *xanA* in *X*. *citri*, encode the same enzyme, phosphomannomutase/phosphoglucomutase (Stevenson *et al*., 1991; Zielinski *et al*., 1991; Köplin *et al*., 1992; McKay *et al*., 2003). Phosphomannomutase (PMM, EC 5.4.2.8) catalyzes reversible interconvertion of α-D-mannose 6-phosphate and α-D-mannose 1-phosphate, a key reaction in the biosynthesis of the colanic acid, alginate, and xanthan gum (Zielinski *et al*., 1991; Köplin *et al*., 1992). This enzyme also functions as phosphoglucomutase (PGM, EC 5.4.2.2), catalyzing interconvertion of α-D-glucose 6-phosphate and α-D-glucose 1-phosphate that is involved in synthesis of the bacterial lipopolysaccharide. In *P*. *aeruginosa*, it is also involved in production of rhamnolipid surfactants (Olvera *et al*., 1999). The same α-D-phosphohexomutase superfamily also includes phosphoglucosamine mutase (GlmM, EC 5.4.2.10) that catalyzes the conversion of α-glucosamine-6-phosphate to α-glucosamine-1-phosphate, which is involved in peptidoglycan and lipopolysaccharide biosynthesis (Mehra-Chaudhary *et al*., 2011a). Bacterial GlmMs reportedly have a PMM activity of about 20% of its phosphoglucosamine mutase activity and a low PGM activity (Tavares *et* al., 2000). The biochemical and structural properties of these enzymes have been extensively characterized, with high-resolution crystal structures available for *P*. *aeruginosa* AlgC (Regni *et al*., 2002; Regni *et al*., 2004), *X*. *citri* XanA (Goto *et al*., 2016), human PGM1 (Stiers and Beamer, 2018) and PMMs from several other organisms, as well as for *Bacillus anthracis* GlmM (Mehra-Chaudhary *et al*., 2011a). The phylogenetic relationships between these enzymes and the structural basis of the enzyme specificity have been described in detail (Whitehouse *et al*., 1998; Regni *et al*., 2004; Shackelford *et al*., 2004), see (Stiers *et al*., 2017) for a comprehensive review. It should be noted that certain eukaryotes, bacteria and archaea encode a distinct form of PMM/PGM, a member of the haloacid dehalogenase (HAD) superfamily (Zhang *et al*., 2018), that has a distinct stuctural fold and therefore represents an analogous (non-homologous isofunctional) enzyme (Omelchenko *et al*., 2010).

While eukaryotic PMM/PGMs have been extensively studied since 1950s, the first bacterial enzymes of this family have been cloned and characterized in 1991 (Jiang *et al*., 1991; Stevenson *et al*., 1991; Zielinski *et al*., 1991). The cloned 463-aa ORF from *P*. *aeruginosa*, which restored mucoid phenotype to an alginate-negative mutant, represented the enzymatically active C-terminal part of the protein, starting from the Met392 of the “long” ORF (Table 1 and Supporting Information Figure S3). All subsequent studies assumed this fragment to be the full-length protein, and its upstream DNA region (GenBank accession L00980), sequenced shortly thereafter (Zielinski *et al*., 1992), has not been recognized as protein-coding. Further, in the process of curation at UniProtKB, the full-length AlgC sequence of *P*. *aeruginosa* strain PA14 (locus tag PA14_70270, GenBank accession ABJ14705.1) has been marked as ‘erroneus initiation’, and the N-terminal periplasmic domain was removed from the ALGC_PSEAB entry (Table 1). The data presented in Supporting Information Figure S5 clearly show that the full-length ORF PA14_70270 of *P*. *aeruginosa* strain PA14 is in fact transcribed and there is no reason to assume that it could not be translated, as has been shown here for the *L*. *enzymogenes* protein Le2152 (Supporting Information Figure S2A).

The combination of a periplasmic dCache-type sensor domain with a cytoplasmic enzymatic domain suggests that the activity of the membrane-anchored PMMs could be modulated by external ligands, such as C4-dicarboxylates or acetate that are sensed by some dCache domains (Cheung and Hendrickson, 2008; Zhou *et al*., 2008; Pineda-Molina *et al*., 2012); the ligand sensed by the closely related *dCache_1* domain of *B*. *subtilis* sporulation kinase KinD remains to be identified. If so, two-domain PMMs would offer yet another example of a sensory system that regulates bacterial metabolism in response to environmental cues (Galperin, 2004, 2018). Environmental regulation of the PMM/PGM activity would seem justified, based on the unique position of this enzyme as a common step in peptidoglycan, lipopolysaccharide, and exopolysaccharide biosynthesis pathways. In *P*. *aeruginosa*, AlgC has been even referred to as a checkpoint enzyme that coordinates biosynthesis of alginate, Pel, Psl and lipopolysaccharide (Ma *et al*., 2012). However, regulation by dCache domains is typically mediated by their dimerization, which in turn leads to dimerization and, hence, activation of the downstream enzymatic or methyl-acceptor domains (Ortega *et al*., 2017). All available data indicate that PMMs are active as monomers, although some PGMs and GlmMs have been seen to form dimers (Mehra-Chaudhary *et al*., 2011a; Mehra-Chaudhary *et al*., 2011b; Stiers *et al*., 2017). Future experiments will be needed to figure out the mechanisms of regulation of AlgC-type PMMs. It is quite likely that the periplasmic dCache domains inhibit, rather than activate, the PMM activity of the respective cytoplasmic domains. It is important to note that most data obtained by studying (short) AlgC enzymes would not be affected by the sequence correction suggested in this work. Deletion of the cytoplasmic domain of a long PMM abolishes its enzymatic activity, so all mutation data remain valid. Further, the RNA-Seq data examined in the course of this work (Supporting Information Figure S5) suggest the existence of at least two transcription starts, so there is a distinct possibility that a “short” ORF could be expressed *in vivo* from the “long” AlgC template. Therefore, *algC* expression and its PMM/PGM activity are likely to be regulated at both the transcriptional and post-transcriptional level. *Lysobacter enzymogenes*, where AlgC is directly involved in production of type IV pili, could serve as an attractive model to untangle the complex regulation of the PMM/PGM activity and its role in various cellular processes.

On a more general note, this Genome Update shows the value of examining the genomic data even for very well characterized enzymes.

## EXPERIMENTAL PROCEDURES

### Bacterial strains, plasmids and growth conditions

The bacterial strains and plasmids used in this study are listed in Supporting Information Table S1. Unless stated otherwise, *L*. *enzymogenes* was grown in LB medium or 1/10 Tryptic Soy Broth (TSB) at 28 °C with appropriate antibiotics -- kanamycin (Km), 25 μg/mL, for mutant construction and gentamicin (Gm), 150 μg/mL, for plasmid maintenance.

### Genetic methods

Double-crossover homologous recombination was used to generate mutants in *L*. *enzymogenes* OH11, as described previously (Qian *et al*., 2012), using primers listed in Supporting Information Table S2. In brief, two flanking regions of *Le2152* were generated by PCR amplification and cloned into the suicide vector pEX18Gm (Supporting Information Table S1). The final constructs were transformed into the wild-type strain by electroporation. The single-crossover recombinants were selected on LB plates supplemented with Km and Gm. The recombinants were cultured in LB without antibiotics for 6 h and subsequently plated on LB agar containing 10% (w/v) sucrose and Km. The sucrose-resistant, Km-resistant but Gm-sensitive colonies representing double crossovers were picked up. In-frame gene deletions were verified by PCR using appropriate primers (Supporting Information Table S2).

Gene complementation construct was generated as described earlier (Qian *et al*., 2014). Briefly, the DNA fragment containing the coding region of Le2152 and its native promoter region was amplified by PCR with designated primer pairs (Supporting Information Table S2) and cloned into the broad-host vector pBBR1-MCS5 (Supporting Information Table S1). The plasmid was transformed into the mutant by electroporation, and the transformants were selected on the LB plates containing Km and Gm.

### Twitching motility assay

Twitching motility of *L*. *enzymogenes* OH11 was assayed as described previously (Wang *et al*., 2014). In general, a thin layer of 1 mL 1/20 tryptic soy agar medium supplemented with 1.8% agar was evenly spread onto a sterilized microscope slide. To create a thin inoculation line, the edge of a sterilized coverslip was dipped into the bacterial cell suspension and then gently pressed onto the surface of the medium. After 24 h incubation, the margin of the bacterial culture on the microscope slide was observed under a microscope at 640X magnification. Twitching motility of *L*. *enzymogenes* was indicated by the presence of individual mobile cells or small clusters of cells growing outwardly from the main colony, as described in our earlier report (Wang *et al*., 2014). Three replicate slides were used for each sample, with the experiment carried out three times.

### Sequence analysis

Iterative sequence similarity searches were performed using PSI-BLAST (Altschul *et* al., 1997) and jackHMMer (Potter *et al*., 2018). The transmembrane orientation and domain architectures of long PMMs were predicted using TMHMM (Krogh *et al*., 2001) and verified by checking the InterPro (Mitchell *et al*., 2019) entries, where available. Genomic neighborhoods of “long” and “short” PMM genes were examined using the SEED (Overbeek *et al*., 2014) and the NCBI Genome database and plotted using the respective genomic coordinates. Alignments of the predicted periplasmic domains and the flexible linkers were generated from jackHMMer outputs and used to create the sequence logos with WebLogo (Crooks *et al*., 2004). Phylogenetic analysis of the enzymatic domains of short and long PMMs was peformed with MEGA7 (Kumar *et al*., 2016) using an alignment generated by MUSCLE (Edgar, 2004) and manually trimmed to remove non-enzymatic domains.

Analysis of the RNA-Seq data was performed by searching the NCBI’s Sequence Read Archive (SRA) with MegaBLAST (Zhang *et al*., 2000). Genomic fragments coding for the long PMM genes from Table 1 were expanded to 3 kb in such a way that each of them included the *algC* upstream region and a part of the *dut* ORF. These 3-kb DNA fragments were used to query the SRA entries for the respective organisms using discontiguous MegaBLAST with default parameters, except that ‘Max target sequences’ number was increased to 1000. The results from each MegaBLAST search were examined by setting the ‘Graphical overview’ parameter to 1000 sequences and checking for hits that correspond to the N-terminal fragments of the “long” PMMs.

## ACKNOWLEDGEMENTS

This work was supported by the National Natural Science Foundation of China (31872016 to GQ), the Fundamental Research Funds for the Central Universities (KYT201805, Y0201600126 and KYTZ201403 to GQ), Natural Science Foundation of Jiangsu Province (BK20181325 to GQ), Innovation Team Program for Jiangsu Universities (2017 to GQ) and by the NIH Intramural Research Program at the National Library of Medicine (MYG).

## Supporting Information

Table S1. Strains and plasmids used in this study

Table S2. Primers used in this study

Table S3. Phosphomannomutases and phosphoglucosamine mutases encoded in selected gammaproteobacterial genomes

Table S4. Identification of two-domain phosphomannomutase transcripts in RNA-seq samples

Figure S1. Twitching motility of *Lysobacter enzymogenes* is not affected by the absence of the *glmM* gene.

Figure S2. Molecular weight and domain organization of the Le2152 protein.

Figure S3. Genomic neighborhoods of the *Xanthomonas* spp. and *Pseudomonas* spp. “long” PMM genes according to the SEED database.

Figure S4. Maximum likelihood phylogenetic tree of the cytoplasmic enzymatic domains of “long” and “short” phosphomannomutases.

Figure S5. RNA-Seq data for the “long” PMMs from *Lysobacter enzymogenes* and *Pseudomonas aeruginosa*.

