## Supplemental Tables S1-S4 and Supplemental Figures S1-S5 for "Two forms of phosphomannomutase in gammaproteobacteria: The overlooked membrane-bound form of AlgC is required for twitching motility of *Lysobacter enzymogenes*"

submitted for publication in *Environmental Microbiology*

**Table S1. Strains and plasmids used in this study**

| **Strains** | **Characteristics**[a] | **Source** |
|---|---|---|
| ***Lysobacter enzymogenes*** | | |
| OH11 | Wild type strain, $Km^R$ | (Qian *et al.*, 2009) CGMCC No. 1978 |
| Δ*Le2152* | In-frame deletion of *Le2152* (*D9T17_01580*) in the background of strain OH11, $Km^R$ | This study |
| Δ*Le2152* (pBBR) | Δ*Le2152* carrying the empty vector pBBR1-MCS5, $Km^R$, $Gm^R$ | This study |
| Δ*Le2152* (*2152*) | Δ*le2152* carrying the pBBR1-Le2152 plasmid, $Km^R$, $Gm^R$ | This study |
| Δ*Le4861* | In-frame deletion of *Le4861 (*D9T17_13225*)* in the background of strain OH11, $Km^R$ | This study |
| ***Escherichia coli*** | | |
| DH5α | Host strain for molecular cloning | |
| **Plasmids** | | |
| pBBR1-MCS5 | Broad-host vector with a $P_{lac}$ promoter, $Gm^R$ | (Kovach *et al.*, 1995) |
| pBBR1-Le2152 | pBBR1-MCS5 cloned with a 3,002-bp fragment (RCTY01000005: 280-3281), in which *Le2152* was under its native promoter, $Gm^R$ | This study |

[a]$Km^R$, $Gm^R$ = Kanamycin-, Gentamicin-resistance, respectively.

**Table S2. Primers used in this study**

| Primer | Sequence[a] (5'-3')[a] | Use |
|---|---|---|
| Construction of complementation plasmids | | |
| *Le2152*-F | CGGAATTCGATTACCAGGGACCGCTGTT (*Eco*RI) | To amplify the fragment of intact *Le2152* sequence |
| *Le2152*-R | GCTCTAGACGCCGAAAACGAAAACGCCG (*Xba*I) | |
| In-frame deletion | | |
| *Le2152* F1 | CGGGATCCGATTACCAGGGACCGCTGTT (*Bam*HI) | To amplify a 465-bp upstream arm of *Le2152* |
| *Le2152* R1 | CCCAAGCTTCTTCACTTCGACCATAACCT (*Hin*dIII) | |
| *Le2152* F2 | CCCAAGCTTGCCGATCCTGGTGATGCGTT (*Hin*dIII) | To amplify a 654-bp downstream arm of *Le2152* |
| *Le2152* R2 | GCTCTAGATGGACGAAACGGTCAAGGCC (*Xba*I) | |

[a]Underlined nucleotide sequence are restriction sites and the restriction enzymes are indicated

**Table S3. Phosphoglucosamine mutase and two types of phosphomannomutase encoded in selected bacterial genomes**

| Organism | Locus tag (GenBank accession number) | | |
|---|---|---|---|
| | PMM short version (XanA) | PMM long version (AlgC) | GlmM |
| Xanthomonadales | | | |
| *Xanthomonadaceae* | | | |
| *Lysobacter enzymogenes* OH11 | – | D9T17_01580 (ROU09072.2) | D9T17_13225 (ROU06416.1) |
| *Lysobacter enzymogenes* C3 | – | GLE_0254 (ALN55613.1) | GLE_2150 (ALN57500.1) |
| *Lysobacter enzymogenes* M497-1 | – | LEN_4688 (BAW00176.1) | LEN_2888 (BAV98375.1) |
| *Lysobacter antibioticus* 76 | – | LA76x_4631 (ALN82734.1) | LA76x_2994 (ALN81122.1) |
| *Lysobacter* sp. TY2-98 | – | DWG18_12515 (AXK73019.1) | DWG18_06235 (AXK71922.1) |
| *Luteimonas* sp. JM171 | – | BGP89_04110 (AOH35641.1) | BGP89_11115 (AOH36832.1) |
| *Luteimonas* sp. 100111 | CNR27_14690 (ATD68527.1) | CNR27_02850 (ATD66522.1) | CNR27_06990 (ATD67218.1) |
| *Pseudoxanthomonas suwonensis* 11-1 | Psesu_2445 (ADV28277.1) | Psesu_0157 (ADV26019.1) | Psesu_1849 (ADV27691.1) |
| *Stenotrophomonas maltophilia* K279a | Smlt0653 (CAQ44236.1) | Smlt0403 (CAQ44002.1) | Smlt3414 CAQ46841.1) |
| *Xanthomonas campestris* pv. *campestris* str. ATCC 33913 | XCC0626 (AAM39942.1, P0C7J2) | XCC3857* (AAM43088.1*, Q8P459*)[a] | XCC2539(AAM41811.1, Q8P7S2) |
| *Xanthomonas citri* pv. *citri* str. 306 | XAC3579 (AAM38422.1, Q8PGN7, PDB: 5BMN) | XAC3912* (AAM38749.1*, Q8PFR6*)[b] | XAC2714 (AAM37559.1, Q8PJ31) |
| *Xanthomonas citri* subsp. *citri* Aw12879 | XCAW_04279 (AGI10044.1) | XCAW_00367 (AGI06191.1) | XCAW_01461 (AGI07263.1) |
| *Xanthomonas citri* pv. *malvacearum* XcmN1003 | APY30_03860 (ASN08225.1) | APY30_02215 (ASN07911.1) | APY30_13635 (ASN10039.1) |
| *Xanthomonas arboricola* pv. *juglandis* Xaj 417 | AKJ12_15195 (AKU50990.1) | AKJ12_21745 (AKU52111) | AKJ12_03610 (AKU48962.1) |
| *Xanthomonas axonopodis* Xac29-1 | XAC29_18220 (AGH79049.1) | XAC29_19845 (AGH79364.1) | XAC29_13835 (AGH78203.1) |
| *Xanthomonas oryzae* pv. *oryzae* PXO99A | PXO_03174 (ACD61040.1) | PXO_02922 (ACD61293.1) | PXO_01274 (ACD60107.1) |
| *Xanthomonas translucens* pv. *undulosa* Xtu 4699 | FD63_16520 (AKK68964.1) | FD63_01870 (AKK66326.1) | FD63_06880 (AKK67226.1) |
| *Xylella fastidiosa* 9a5c | XF_0260 (AAF83073.1) | XF_0151* (AAF82964.1*)[c] | XF_1468 (AAF84277.1) |
| *Xylella taiwanensis* PLS235 | AB672_01195 (AXI82683.1) | AB672_00665 (AXI82593.1) | AB672_08140 (AXI83904.1) |

| | | | |
|---|---|---|---|
| **Rhodanobacteraceae** | | | |
| *Ahniella affigens* D13 | – | C7S18_20435 (AVQ00208.1) | C7S18_10840 (AVP97666.1) |
| *Dokdonella koreensis* DS-123 | – | I596_3660 (ANB19648.1) | I596_1334 (ANB17364.1) |
| *Dyella japonica* A8 | HY57_20075 (AIF49391.1) | HY57_02855 (AIF46264.1) | HY57_11910 (AIF47916.1) |
| *Frateuria aurantia* DSM 6220 | Fraau_2860 (AFC87194.1) Fraau_3262 (AFC87585.1)[d] | – | Fraau_1995 (AFC86380.1) |
| *Luteibacter rhizovicinus* DSM 16549 | BJI69_11355 (APG06496.1) | BJI69_08495 (APG03940.1) | BJI69_20240 (APG05999.1) |
| *Rhodanobacter denitrificans* 2APBS1 | R2APBS1_0585 (AGG87754) | R2APBS1_3851 (AGG90903) | R2APBS1_1877 (AGG89001), R2APBS1_2133 (AGG89250) |
| **Pseudomonadales** | | | |
| *Pseudomonas aeruginosa* PAO1 | – | PA5322* (AAG08707.1*, P26276*, WP_003121305*)[e] | PA4749 (AAG08135.1, Q9HV50) |
| *Pseudomonas aeruginosa* UCBPP-PA14 | – | PA14_70270 (ABJ14705.1, Q02E40*)[f] | PA14_62840 (ABJ14132.1, Q02FS3) |
| *Pseudomonas aeruginosa* M1608 | – | HW04_21660 (ALY73569.1) | HW04_06110 (ALY70583.1) |
| *Pseudomonas alcaligenes* strain NEB 585 | – | A0T30_01015 (AMR65011.1) | A0T30_18140 (AMR68200.1) |
| *Pseudomonas brassicacearum* LBUM | AK973_6087(ALQ06536.1) | – | AK973_0978(ALQ01427.1) |
| *Pseudomonas citronellolis* P3B5 | – | PcP3B5_60560 (AMO79415.1) | PcP3B5_54240 (AMO78795.1) |
| *Pseudomonas fluorescens* SBW25 | PFLU_5986 (CAY53511.1)[g] | – | PFLU_5258 (CAY52358.1) |
| *Pseudomonas mendocina* ymp | – | Pmen_4379 (ABP87126.1) | Pmen_3613 (ABP86361.1) |
| *Pseudomonas protegens* Pf-5 | PFL_6054 (AAY95242.1)[h] | – | PFL_0838 (AAY96238.1) |
| *Pseudomonas putida* F1 | Pput_5197 (ABQ81315.1)[i] | – | Pput_4582 (ABQ80702.1) |
| *Pseudomonas resinovorans* NBRC 106553 | – | PCA10_55240 (BAN51256.1) | PCA10_49160 (BAN50648.1) |
| *Pseudomonas stutzeri* A1501 | – | PST_0470 (ABP78176.1) | PST_3317 (ABP80946.1) |
| *Pseudomonas syringae* pv. *tomato* DC3000 | – | PSPTO_0083* (AAO53637.1*, Q88BD4*)[j] | PSPTO_4495 (AAO57943.1, Q87WQ0) |
| **Alteromonadales** | | | |
| *Marinobacter pelagius* 114J | DET50_10559 (RBP31835.1) DET50_105143 (RBP31915.1) | DET50_10255 (RBP33243.1) | DET50_102236 (RBP33422.1) |
| *Marinobacter* sp. CP1 | ACP86_12145 (AKV96849.1) | ACP86_07110 (AKV95939.1) | ACP86_05865 (AKV95727.1) |

| | | | |
|---|---|---|---|
| **Cellvibrionales** | | | |
| *Cellvibrio japonicus* Ueda107 | CJA_2117 (ACE83813.1) | CJA_3524 (ACE86337.1) | CJA_2672 (ACE83954.1) |
| *Congregibacter litoralis* KT71 | KT71_00920 (EAQ98495.1) | KT71_05085 (EAQ97656.1) | KT71_07979 (EAQ97302.1) |
| *Kangiella koreensis* DSM 16069 | – | Kkor_2220 (ACV27629.1) | Kkor_2365 (ACV27774.1) |
| *Microbulbifer* sp. A4B17 | BTJ40_02900 (AWF79849.1) | BTJ40_21625 (AWF83206.1) | BTJ40_18190 (AWF82580.1) |
| *Pseudohaliea rubra* DSM 19751 | HRUBRA_02500 (KGE02964.1) | HRUBRA_01211 (KGE04182.1) | HRUBRA_02668 (KGE02690.1) |
| *Saccharophagus degradans* 2-40 | Sde_1001 (ABD80263.1) | Sde_3676 (ABD82931.1) | Sde_2718 (ABD81978.1) |
| *Teredinibacter turnerae* T7901 | TERTU_0950 (ACR11266.1) | TERTU_0186 (ACR13976.1) | TERTU_3262 (ACR12878.1) |
| **Chromatiales** | | | |
| *Nitrococcus mobilis* Nb-231 | NB231_04975 (EAR22234.1) | NB231_11214 (EAR20842.1) | NB231_03862 (EAR20881.1) |
| *Nitrosococcus halophilus* Nc 4 | – | Nhal_3802 (ADE16817.1) | Nhal_3703 (ADE16723.1) |
| *Nitrosococcus watsonii* C-113 | – | Nwat_3050 (ADJ29771.1) | Nwat_0553 (ADJ27516.1) |
| *Thioflavicoccus mobilis* 8321 | Thimo_0977 (AGA89803.1) | Thimo_0741 (AGA89580.1) | Thimo_2546 (AGA91270.1) |
| **Methylococcales** | | | |
| *Methylobacter tundripaludum* SV96 | Mettu_4200 (EGW21045.1) | Mettu_1833 (EGW22995.1) | Mettu_3203 (EGW20077.1) |
| *Methylococcus capsulatus* str. Bath | – | MCA2782* (AAU91112.1*)[k] | MCA1847 (AAU91923.1) |
| *Methylomicrobium alcaliphilum* 20Z | MEALZ_3227 (CCE24892.1) | MEALZ_0132 (CCE21833.1) | MEALZ_1810 (CCE23496.1) |
| *Methylomonas methanica* MC09 | Metme_3866 (AEG02220.1) | Metme_3325 (AEG01696.1) | Metme_3291 (AEG01662.1) |
| **Oceanospirillales** | | | |
| *Alcanivorax borkumensis* SK2 | ABO_0937 (CAL16385.1) | ABO_0211 (CAL15659.1) | ABO_0324 (CAL15772.1) |
| *Alcanivorax pacificus* W11-5 | S7S_10425 (AJD48497.1) | S7S_00400 (AJD46504.1) | S7S_16515 (AJD49715.1) |
| *Gynuella sunshinyii* YC6258 | YC6258_03897 (AJQ95933.1) | YC6258_01082 (AJQ93130.1) | YC6258_05251 (AJQ97281.1) |
| *Hahella chejuensis* KCTC 2396 | HCH_06439 (ABC33084.1) | HCH_01025 (ABC27908.1) | HCH_01234 (ABC28106.1) |
| *Oleispira antarctica* RB-8 | OLEAN_C25180 (CCK76694) | OLEAN_C38520 (CCK78028) | OLEAN_C32860 (CCK77462) |
| *Thalassolituus oleivorans* MIL-1 | TOL_2551 (CCU72950.1) | TOL_0208 (CCU70656.1) | TOL_2722 (CCU73119.1) |
| **Thiotrichales** | | | |
| *Cycloclasticus* sp. PY97N | – | CPC19_03270 (ATI02519.1) | CPC19_05560 (ATI02953.1) |
| *Cycloclasticus zancles* 78-ME | CYCME_0971 (AGS39304.1) | CYCME_0529 (AGS38870.1) | CYCME_1015 (AGS39348.1) |

* Asterisks indicate long (>700 aa) phosphomannomutase sequences that are listed in GenBank and/or UniProt databases in an N-terminally truncated form.

[a] - Corrected 782-aa sequence of XCC3857 has been submitted to RefSeq.

[b] - Corrected 781-aa sequence of XAC3912 is available as RefSeq entry WP_015471480.

[c] - Corrected 787-aa sequence of XF_0151 is available as RefSeq entry WP_023906061.

[d] - The gap between *dut* and *algC* (Fraau_3262) genes is 148 bp and contains three stop codons.

[e] - UniProt entry P26276 lacks 405 N-terminal amino acid residues, the full length 868-aa ORF is the RefSeq entry NP_254009.2.

[f] - UniProt entry Q02E40 lacks 391 N-terminal amino acid residues, removed as ‘erroneus initiation’. ABJ14705.1 is the full-length entry.

[g] - The gap between *dut* and *algC* genes is 265 bp and contains two stop codons.

[h] - The gap between *dut* and *algC* genes is 262 bp and contains two stop codons.

[i] - The gap between *dut* and *algC* genes is 112 bp and contains three stop codons.

[j] - UniProt entry Q88BD4 lacks 359 N-terminal amino acid residues; the full length 824-aa ORF is equivalent to residues 38-861 of KPY97009.1.

[k] - The sequence of MCA2782 is at least 787 aa long.

**Table S4. Identification of two-domain phosphomutase transcripts in RNA-seq samples**

| Organism used for RNA-Seq | SRA entries | Data reference |
|---|---|---|
| *Lysobacter enzymogenes* OH11 | SRX4451783- SRX4451785, SRX4941505-SRX4941508 | N/A |
| *Stenotrophomonas maltophilia* K279a | | (Abda *et al.*, 2015) |
| *Stenotrophomonas maltophilia* FLR | SRX4893298-SRX4893300 | (Gallagher *et al.*, 2019) |
| *Xanthomonas campestris* pv. *campestris* B100 | SRX3764896 - SRX3764900, ERX1966468 | (Alkhateeb *et al.*, 2017; Alkhateeb *et al.*, 2018) |
| *Xanthomonas campestris* pv. *campestris* 8004 | SRX3312095 - SRX3312097 | (Wang *et al.*, 2017) |
| | ERX1617955 - ERX1617960 | (Bonomi *et al.*, 2016) |
| *Xanthomonas citri* subsp. *citri* Aw12879 | SRX193641, SRX193643, SRX193644 | (Jalan *et al.*, 2013) |
| *Xanthomonas axonopodis* Xac29-1 | SRX4300215, SRX4300216 | N/A |
| *Xanthomonas oryzae* pv. *oryzae* KACC 10331 | SRX235001, SRX235002, SRX707654, SRX707661 | (Kim *et al.*, 2016) |
| *Xylella fastidiosa* subsp. *fastidiosa* 'Temecula' | SRX1286457, SRX1286580, SRX1286607, SRX1286525 | (Parker *et al.*, 2016) |
| *Pseudomonas aeruginosa* PAO1[a] | SRX157659, SRX157683, SRX2792488, SRX4774141, SRX4774147, SRX4804439, SRX591664 | (Cabot *et al.*, 2014; Gill *et al.*, 2018) |
| *Pseudomonas aeruginosa* UCBPP-PA14 | SRX2549303, SRX2549304 | (Deng *et al.*, 2017) |
| *Pseudomonas mendocina* NK-01 | SRX5245299 | (Zhao *et al.*, 2019) |
| *Pseudomonas stutzeri* RCH2 | SRX2990979 - SRX2990984 | (Garber *et al.*, 2018) |
| *Pseudomonas syringae* pv. *tomato* DC3000 | ERX2322626, SRX3984000, SRX3984002, SRX3983989, SRX3644673, SRX3984084 | (Nobori *et al.*, 2018; Schenke *et al.*, 2019) https://gold.jgi.doe.gov/studies?id=Gs0128794 |
| *Cellvibrio japonicus* Ueda107 | SRX2400087, SRX2400089, SRX3542596, SRX3596465 | (Blake *et al.*, 2018) |
| *Methylobacter tundripaludum* 31/32 | SRX2033431, SRX2033432 | (Krause *et al.*, 2017) |
| *Methylococcus capsulatus* str. Bath | SRX2095984, SRX2095985, SRX2095986 | https://gold.jgi.doe.gov/organism?id=Go0000544 |

[a] - Several RNA-seq samples of *P. aeruginosa* PAO1 (e.g. SRX3169723, SRX5457607, SRX5457608) showed expression of the cytoplasmic fragment of AlgC with little or no expression of the N-terminal fragment.

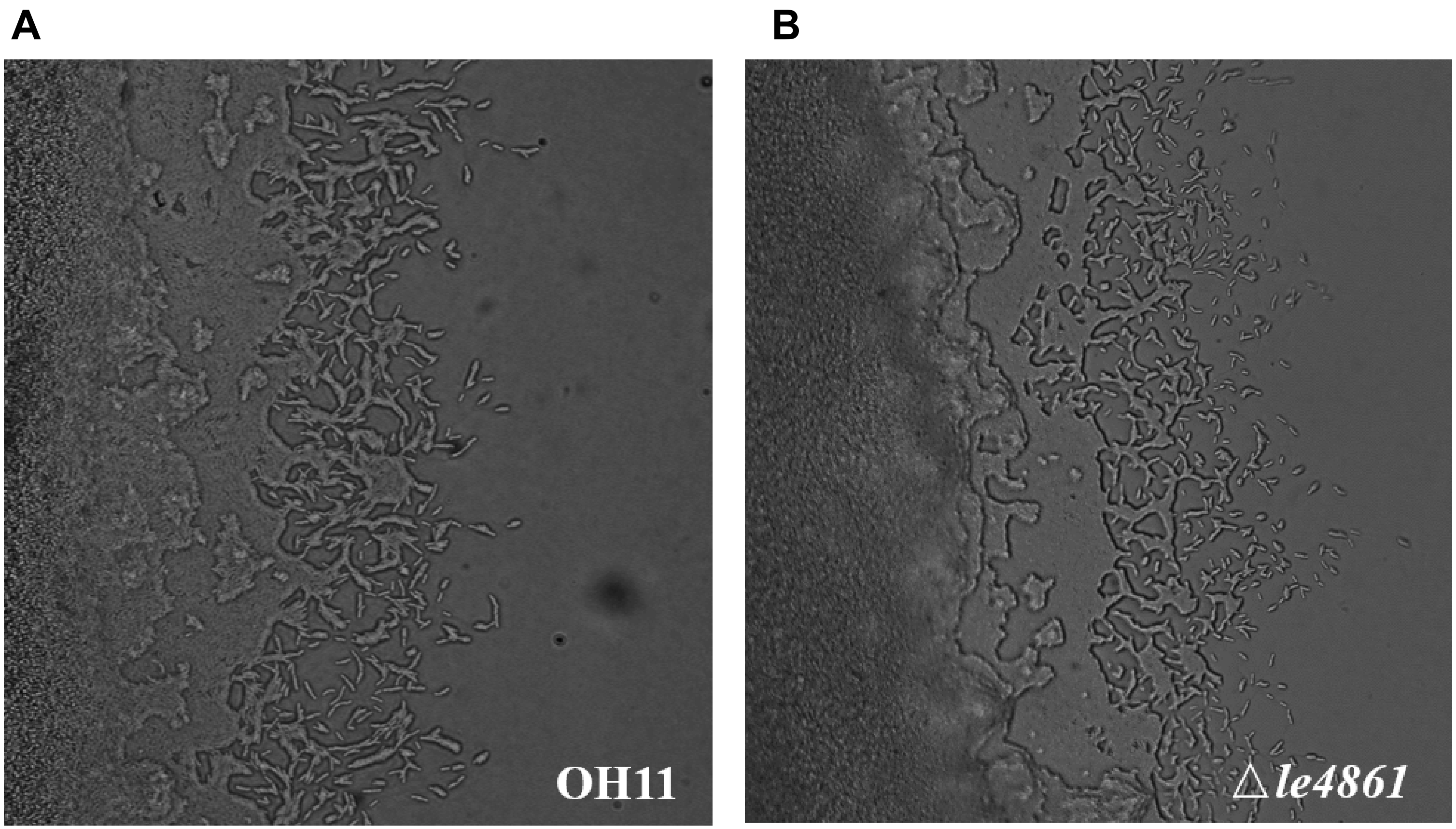


**Figure S1. Twitching motility of *Lysobacter enzymogenes* is not affected by the absence of the *glmM* gene.**

**A.** OH11, wild-type strain of *L. enzymogenes.* **B.** Δ*Le4861*, in-frame deletion mutant of OH11 lacking the phosphoglucosamine mutase gene *Le4861* (locus tag D9T17_13225, GenBank accession ROU06416.1).

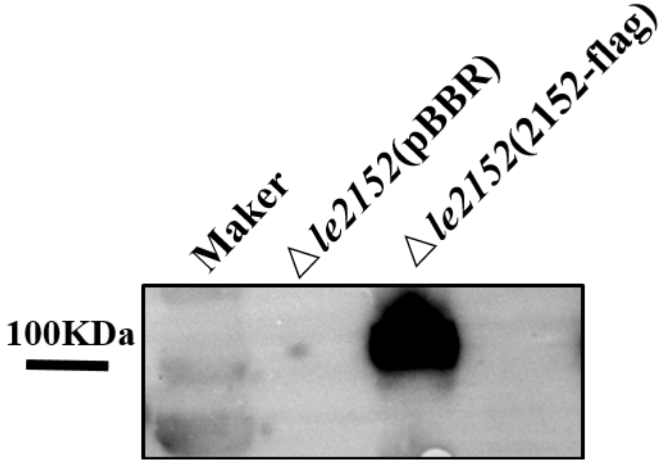


**Figure S2A**. **Expression of the FLAG-labelled Le2152 protein of *Lysobacter enzymogenes*.**
A 3,002-bp DNA fragment containing the Le2152 coding sequence along with its upstream region was cloned into the broad-host vector pBBR1-MCS5. Le2152 was expressed from its native promoter as a FLAG-tagged protein (indicated as 2152-flag) in the *L. enzymogenes Le2152* deletion mutant (Δ*le2152*), grown in the twitching motility-producing medium (10% TSB) to $OD_{600}$ of 1.5. The same strain with an empty vector, indicated as Δ*le2152*(pBBR), was used as control. The blot was probed with anti-FLAG antibody (Abcam, Cambridge, UK). The bar indicates the position of the 100-kDa molecular weight marker (ThermoFisher, Waltham, USA).

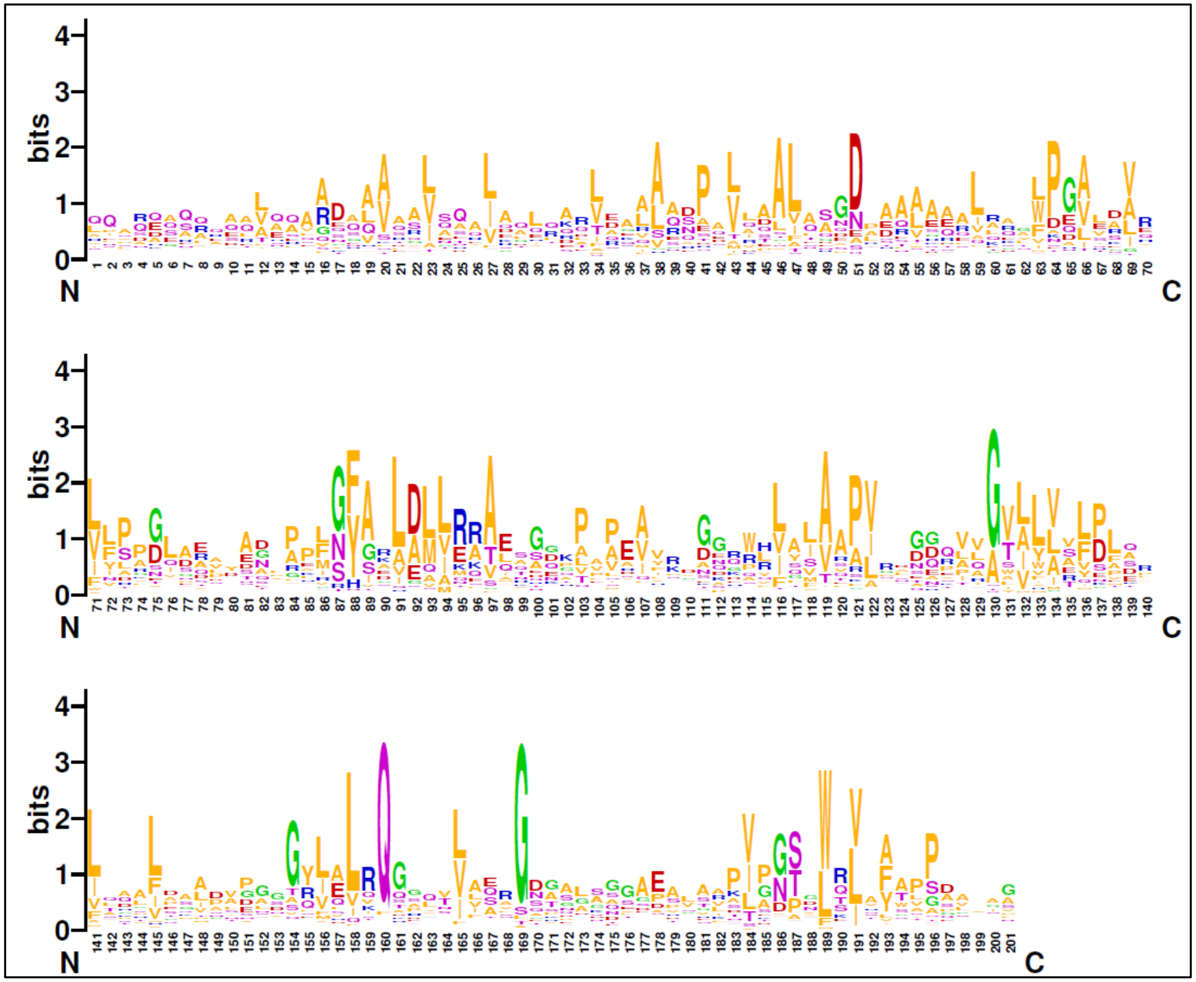


**Figure S2B**. **Sequence conservation in the N-terminal periplasmic domain of membrane-bound phosphomannomutases.** Sequence logo was generated using the WebLogo program, http://weblogo.berkeley.edu (Crooks *et al.*, 2004), from an alignment produced by searching the UniProt database using jackHMMer (Potter *et al.*, 2018) with the N-terminal portion of the Le2152 sequence as the query. Position 1 in the logo corresponds to the Gln40 of Le2152 sequence. The residues are colored as follows: acidic – red; positively charged – blue, hydrophobic – orange; NQST – purple, Gly – green.

```
                 |---------|---------|---------|---------|---------|----------|---------|------
Le2152     40    QQHLDASRRADLIQVRDAAVSSAHDALGAEHKRLRDRLESAPMQAALQAGD-FKAASAELGKDWPGASAGTVLPPDLA   116 (772)
4JGP        7    KDTIAAEHKQEASVLLNLHRNKINYLIGETMARMTSLSIAIDRPVDIKK---MQSILEKTFDSEPRFSGLYFLNAKGD    81 (217)
3BY9        1    MRFQYQALLNEHQSQLDRFSSHIVATLDKYAHIPHLISKDKELVDALLSAQ7TNRYLEQVNEVIQ-AADTYLIDRFGN    83 (260)
3E4P       43    RSQALAGLAGQSRIDASLKASLLRAVVERQRALPLVLADDAAIRGALLSPD6INRKLEALATSAE-AAVIYLIDRSGV   124 (305)
PF14827     1    ----NNMINGQFKQALNMAENHIELIEQMNKTLVNNLAEDHELINYIVTKN3LKDLIVDKRYDIQ-CDQIILLDNKAN    75 (233)
ss_pred          CcchhHHHHHHHHHHHHHHHHHHHHHHHHHHHHHHHHhchhhHHHHhcCC-HHHHHHHHHhhCCCcccEEEEECcChh
ss_dssp          CHHHHHHHHHHHHHHHHHHHHHHHHHHHHHHHHHHHHHHSCSSCCHHH---HHHHHHHHHTTCTTEEEEEEEETTCB

                 ---|------------|---------|----------|---------|----------|---------|---------
Le2152    117    EEYAAL---PKSGYGRLGAMEAAIVADKPMAAIVRD-GGKSVLAMAAPARVG-EQLVGVAYVQLPLQRVGDGVEKADL   189 (772)
4JGP       82    VTASTTELKTKVNLADRSFFIKAKETKKTVISDSYS3TGQPIFTICVPVLDSKRNVTDYLVAAIQIDYLKNLINLLSP   161 (217)
3BY9       84    TIASSNWN5IGRNFAWRPYFYLSIAGQKSQYFALGSTSGQRGYYYAYPVIYA-AEILGVIVVKMDLSAIEQGWQN---   161 (260)
3E4P      125    AVAASNWQ5VGNDYAFRDYFRLAVRDGMAEHFAMGTVSKRPGLYISRRVDGP-GGPLGVIVAKLEFDGVEADWQA---   202 (305)
PF14827    76    VITQSGSAFDGEKLQNLEIVEKTLNQHTSFTTIIRQ-FDIFVLYASAPIVIA-NKQNGMVLIGFSINN--NMMKNIKK   150 (233)
ss_pred          HHHHhC---CCCChhHHHHHHHHHHcCCCceEEEee-CCceEEEEEEeceeC-CEEEEEEEEEechHHhHHHHhhccC
ss_dssp          EEEESSCCSSCCBCTTSHHHHHHHHHCSCEECCCEE-TSSEEEEEEEEEECTTSCEEEEEEEEEEEHHHHHHHHHHHCT

                 |---------|----------|-----------|---------|--------
Le2152    190    GSDSYLALRQG-GFSVIERGDTALN--NGAEALAAKIPGSDLRVAAAVPDIA   238 (772)
4JGP      162    --DVYIEVVNQDGK-MIFASGQASH-AEDQKPVSGYLDDISWNMKVYPNPVT   209 (217)
3BY9      162    -KSSYFVATDDHQV-VFMSSQPAWL51DDYIVSSRPLPELALTIRVLSPKIE   260 (260)
3E4P      203    -SGKPAYVTDRRGIVLITSLPSwrf52AAFLRVETMVPSTNWRLEQLSPLn-   302 (305)
PF14827   151    DTVMDFTVIGDRTVAATS----------------------------------   168 (233)
ss_pred          CCCceEEEEe-CCeeeEEeccHHhhc--chhhheeecCCCceEEEEEcCCCC
ss_dssp          --TSCEEEECTTCC-EEEEESCGGGG-TTCCCEEEECSSSCCEEEECCCCCC
```

**Figure S2C. Sequence alignment of the periplasmic domain of Le2152** with dCache domains from *B. subtilis* sporulation sensor kinase KinD (PDB: 4JGP), dicarboxylate sensor kinase DctB from *Vibrio cholerae* (PDB: 3BY9) and *Sinorhizobium meliloti* (PDB: 3E4P), and the Pfam entry for *dCache_3* domain (PF14827). The alignment was generated with HHpred (Zimmermann *et al.*, 2018). Conserved residues from the sequence logo (Fig. S2B) are shown in bold and colored the same way; ss_pred indicates predicted secondary structure for the dCache domain of Le2152, ss_dssp is the actual secondary structure of 4JGP.

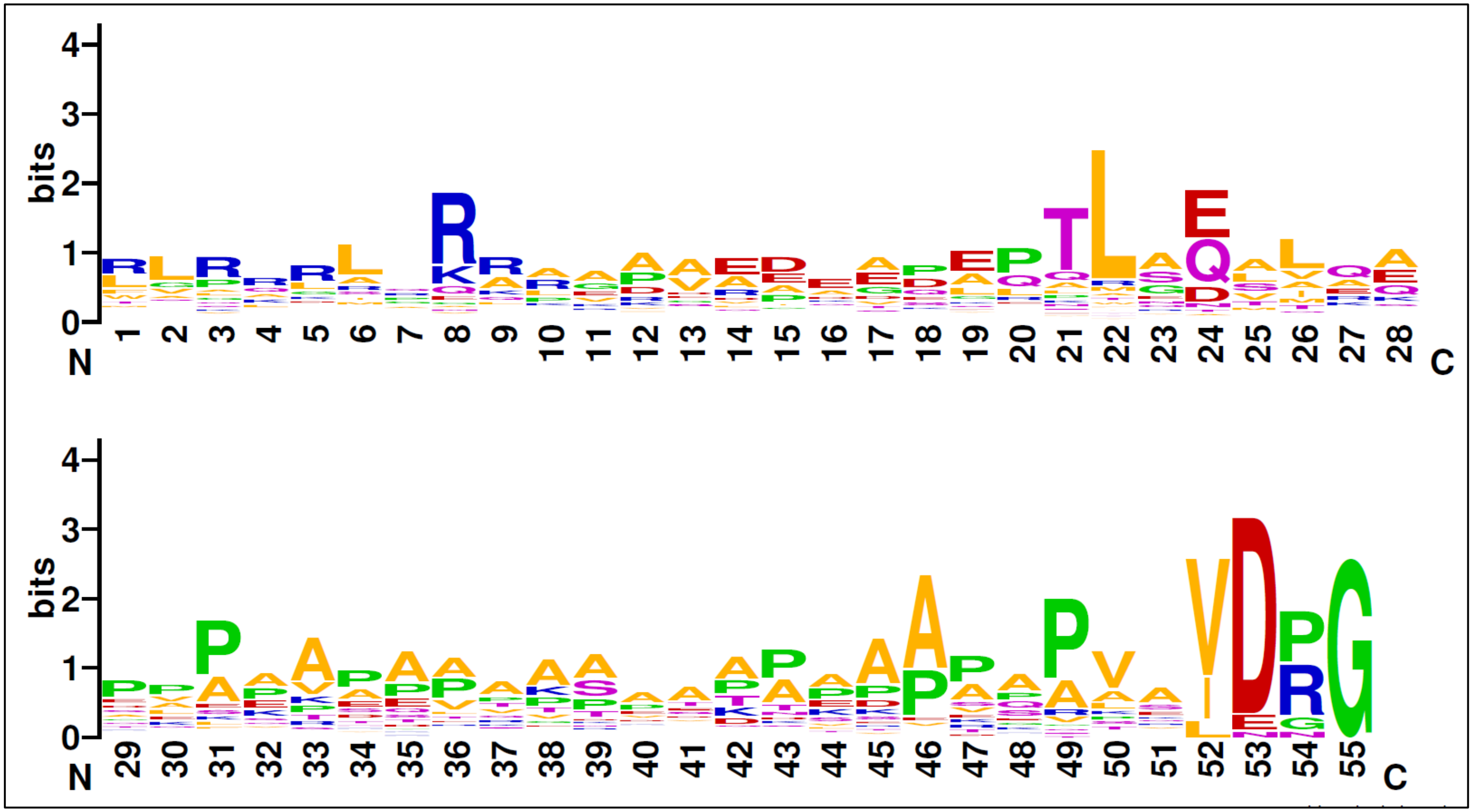


**Figure S2D**. **Sequence conservation in the cytoplasmic linker domain of membrane-bound phosphomannomutases.**
Sequence logo was generated using the WebLogo program, http://weblogo.berkeley.edu (Crooks *et al.*, 2004), from an alignment produced by searching the UniProt database using jackHMMer (Potter *et al.*, 2018) with the N-terminal portion of the Le2152 sequence as the query. Position 1 in the logo correspond to the Arg265 of Le2152 sequence.

```
Le2152      320  IFRAYDIRGVVGQNLDPGIAEMIGQAVGSVMHDQGLSEIVVGRDGRLSGPSLMDGLIAGLRKAGRNVTDIGL  391
1P5D         13  IFRAYDIRGVVGDTLTAETAYWIGRAIGSESLARGEPCVAVGRDGRLSGPELVKQLIQGLVDCGCQVSDVGM   84
5BMN         25  AFKAYDIRGRVPDELNEDLARRIGVALAAQLD---QGPVVLGHDVRLASPALQEALSAGLRASGRDVIDIGL   93
PF02878       3  LFGTSGIRGKVGV2LTPEFALKLGQAIASYLRA2GGGKVVVGRDTRYSSRELARALAAGLAANGVEVILLGL   76

Le2152      392  APTPVTYFGAYHLRAGSCVSLTGSHNPPDYNGFKIVVGGETLSGDAITDLYRRIVGGRLHTAAEPGTLTERD  463
1P5D         85  VPTPVLYYAANVLEGKSGVMLTGSHNPPDYNGFKIVVAGETLANEQIQALRERIEKNDLAS--GVGSVEQVD  154
5BMN         94  CGTEEVYFQTDYLKAAGGVMVTASHNPMDYNGMKLVR3RPISSDTGLFAIRDTVAADTAapGEPTASEQSRT  167
PF02878      77  LPTPALSFATRKLKADGGIMITASHNPPEYNGIKFYD2GGPIPPEVEKKIEAII------------------  131

Le2152      464  ISEDYVQRIASDIQIDRPLQVVVDAGNGVAGDIGPRVLAAIGAEVTPLYCEIDGEFPNHHPDPSEPHNLVDL  535
1P5D        155  ILPRYFKQIRDDIAMAKPMKVVVDCGNGVAGVIAPQLIEALGCSVIPLYCEVDGNFPNHHPDPGKPENLKDL  226
5BMN        168  DKTAYLEHLLSYVDR3KPLKLVVNAGNGGAGLIVDLLAPHLPFEFVRVFHEPDGNFPNGIPNPLLPENRDAT  241
PF02879       1  ---AYIDHLLELVDL5RGLKVVIDPMHGAGGGILPEILKELGAEVVEENCEPDPDFPTRAPNP-EEEALDLL   72

Le2152      536  IKMVQRLNADLGIAFDGDGDRLGVVTRDGQNIFPDRLLMLFAADVLERNPGAMIIYDVKCTGRLPGHILRHGG 608
1P5D        227  IAKVKAENADLGLAFDGDGDRVGVVTNTGTIIYPDRLLMLFAKDVVSRNPGADIIFDVKCTRRLIALISGYGG 299
5BMN        242  AKAVKDNGADFGIAWDGDFDRCFFFDHTGRFIEGYYLVGLLAQAILAKQPGGKVVHDPRLTWNTVEQVEEAGG 314
PF02879      73  IELVKSLGADLGIATDGDADRLGVVDERG-------------------------------------------- 101
PF02880       2  --------------------------------GDQILALLAKYLLEQ4PGAGVVKTVMSSLGLDRVAKKLGG  44

Le2152      609 SPLMWKTGHSLIKAKMRETDAELAGEMSGHFFFKERWYGFDDGIYSAARLLEILAAQPDSPTQTLNALPNGVATPEI 366
1P5D        300 RPVMWKTGHSLIKKKMKETGALLAGEMSGHVFFKERWFGFDDGIYSAARLLEILSQDQRDSEHVFSAFPSDISTPEI 376
5BMN        315 IPVLCKSGHAFIKEKMRSENAVYGGEMSAHHYFRE-FAYADSGMIPWLLIAELVSQSGRSLADLVEAR2KFPCSGEI 391
PF02880      45 KLVRTPVGDKYVKEKMREEGALFGGEESGHIIFLDHAT-TKDGIIAALLVLEILAETGKSLSE-------------- 106

Le2152      367 KVDAPDGDPHSFVERFRNEASFEGARLSTIDGLRVDFPDGWGLVRASNTTPILVMRFDADSQAAMERIQGLFRAQLH 762
1P5D        377 NITVTEDSKFAIIEALQRDAQWGEGNITTLDGVRVDYPKGWGLVRASNTTPVLVLRFEADTEEELERIKTVFRNQLK 453
5BMN        392 NFKV-ADAKASVARVMEhyaslSPELD-YTDGISADFGqWRFNLRSSNTePLLRLNVE3DAALLETRTQ-EISNLLR 467
PF00408       6 ------------VTEKAKVAELAALLKAFADAEKILGEDGRR2VRPSGTEPVLRVMVEADDDELLARLADEVADLLR 71
```

**Figure S2E**. **Sequence alignment of the cytoplasmic enzymatic domain of Le2152** with phosphomannomutase structures from *Pseudomonas aeruginosa* (PDB: 1P5D) and *Xanthomonas citri* (PDB: 5BMN) and with the Pfam domains from PGM_PMM_I to PGM_PMM_IV. Active site residues of experimentally characterized phosphomannomutases (Regni *et al.*, 2004; Goto *et al.*, 2016) are shown in bold and colored as in Fig. S2B.

A

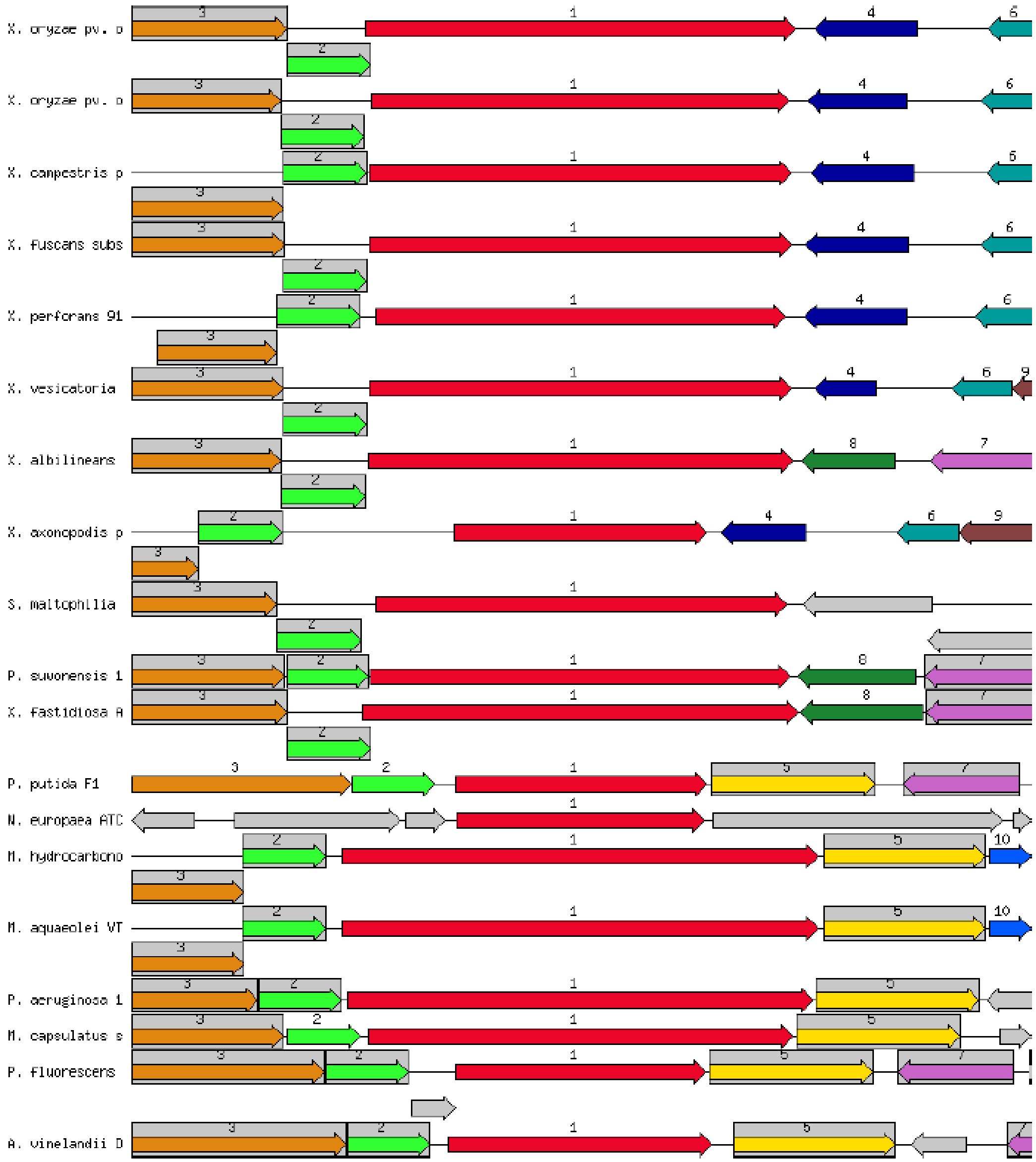

X. oryzae pv. o
X. oryzae pv. o
X. campestris p
X. fuscans subs
X. perforans 91
X. vesicatoria
X. albilineans
X. axonopodis p
S. maltophilia
P. suwonensis 1
X. fastidiosa A
P. putida F1
N. europaea ATC
M. hydrocarbono
M. aquaeolei VT
P. aeruginosa 1
M. capsulatus s
P. fluorescens
A. vinelandii D

B

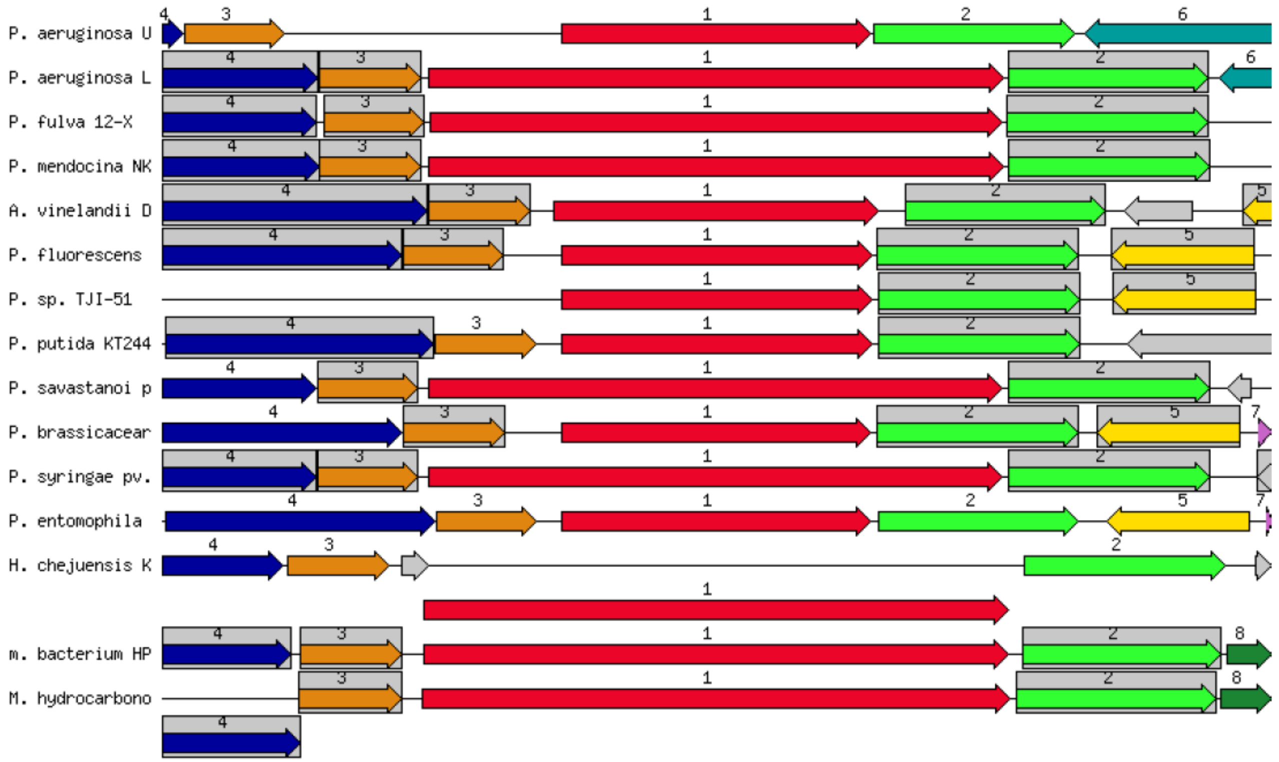


**Figure S3. Genomic neighborhoods of "long" PMM genes according to the SEED database.**

**A.** Homologs of *Xanthomonas oryzae* PXO_02922 gene (GenBank: ACD61293): *algC* genes (marked as 1) are in red, *dut* (no. 2) in green, *coaBC* (no. 3) in orange, *argB* (no. 5) in yellow, uncharacterized gene (no. 4) in dark blue. **B.** Homologs of *Pseudomonas aeruginosa* PA14_70270 gene (GenBank: ABJ14705): *algC* genes (no. 1) are in red, *argB* (no. 2) in green, *dut* (no. 3) in orange, *coaBC* in (no. 4) in dark blue, *pyrE* (no. 5) in yellow. Note the long PMM genes in *X. oryzae*, *X. campestris, X. fuscans,* and other *Xanthomonas* spp., as well as in *S. maltophilia*, *X. fastidiosa* and other organisms (panel A); the short PMM genes in *P. putida*, *P. fluorescens*, and *Azotobacter vinelandii* (panels A and B), and the untranslated N-terminal fragments for the PMMs from *X. axonopodis* (panel A, middle) and *P. aeruginosa* (panel B, top). The data were extracted from the SEED database [http://pubseed.theseed.org/ (Overbeek et al., 2014)] on 16.01.2019.

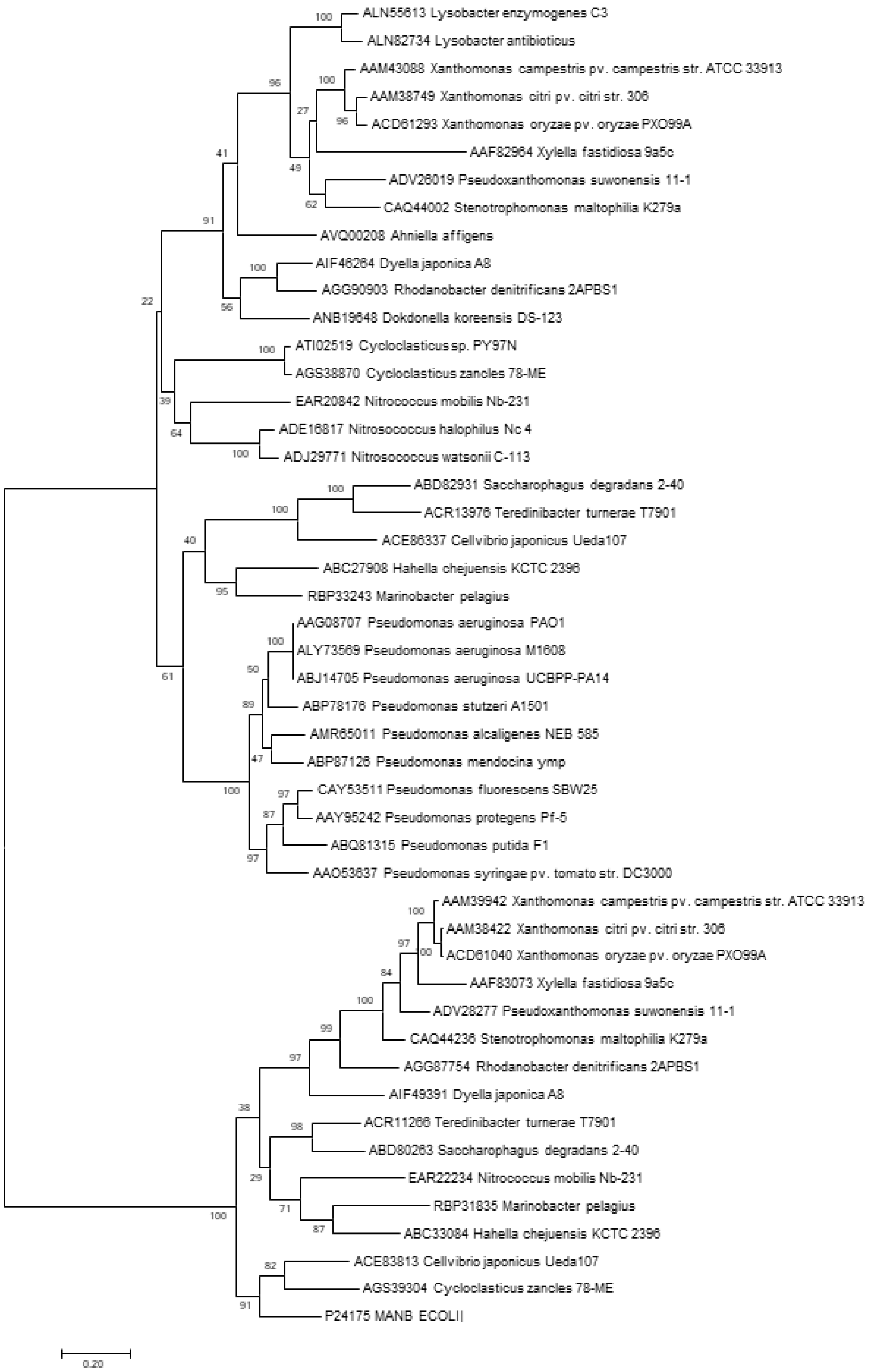


**Figure S4. Maximum likelihood phylogenetic tree of the cytoplasmic enzymatic domains of "long" and "short" phosphomannomutases.** Each protein is listed under its GenBank accession and the respective species name, as in Table S3. The tree was generated with MEGA7 (Kumar *et al.*, 2016) and rooted with the *Escherichia coli* ManB (UniProt: P24175). The bar corresponds to 0.2 substitutions per site.

A

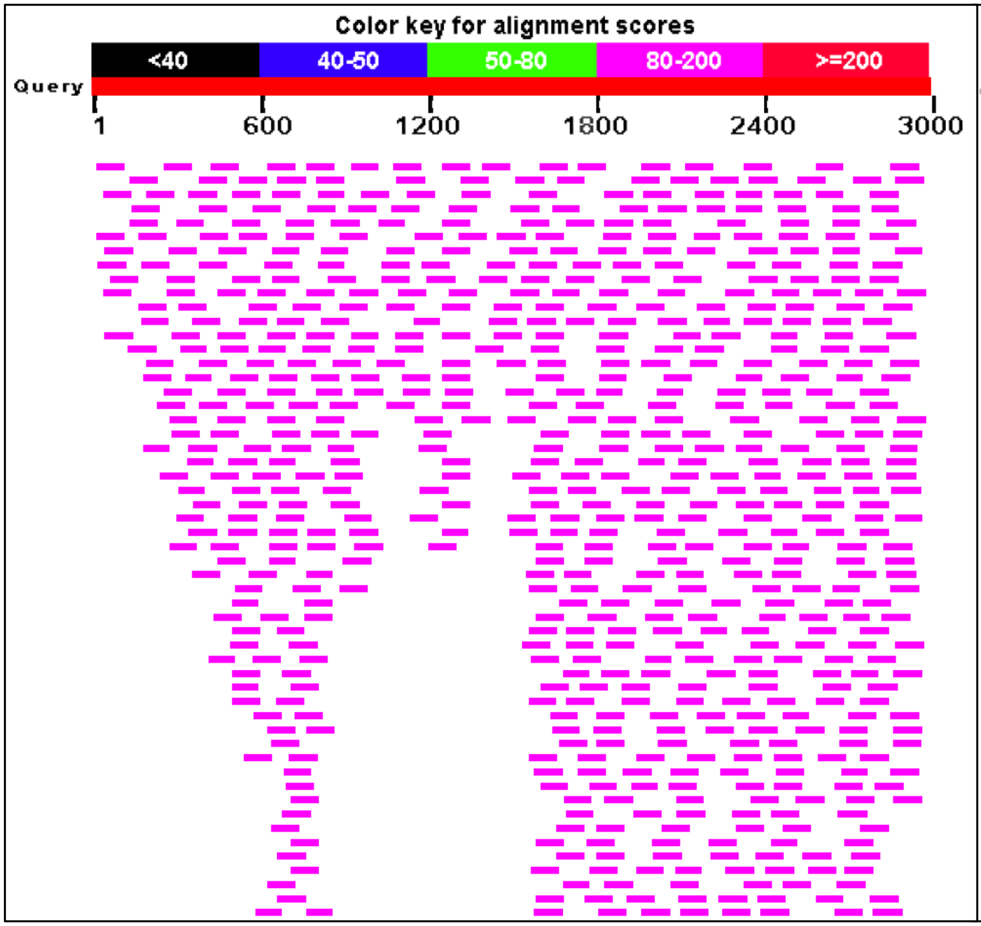
Color key for alignment scores
<40
40-50
50-80
80-200
>=200
Query
1
600
1200
1800
2400
3000

B

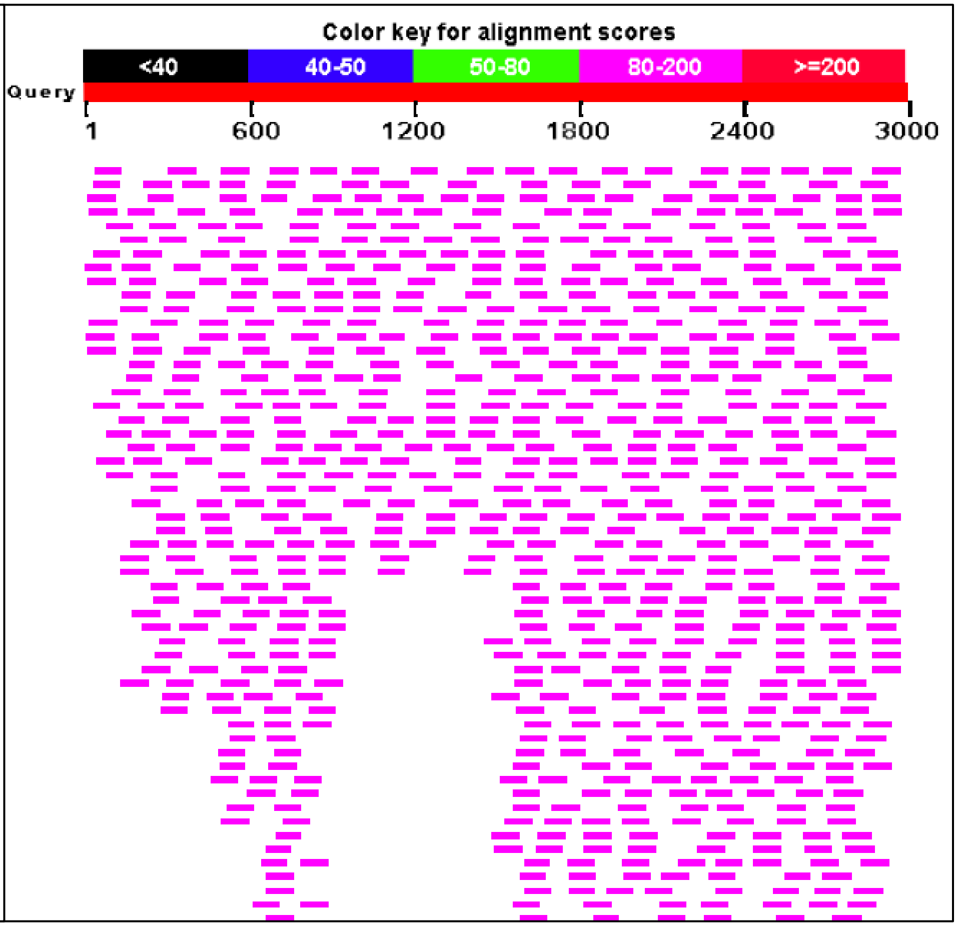
Color key for alignment scores
<40
40-50
50-80
80-200
>=200
Query
1
600
1200
1800
2400
3000

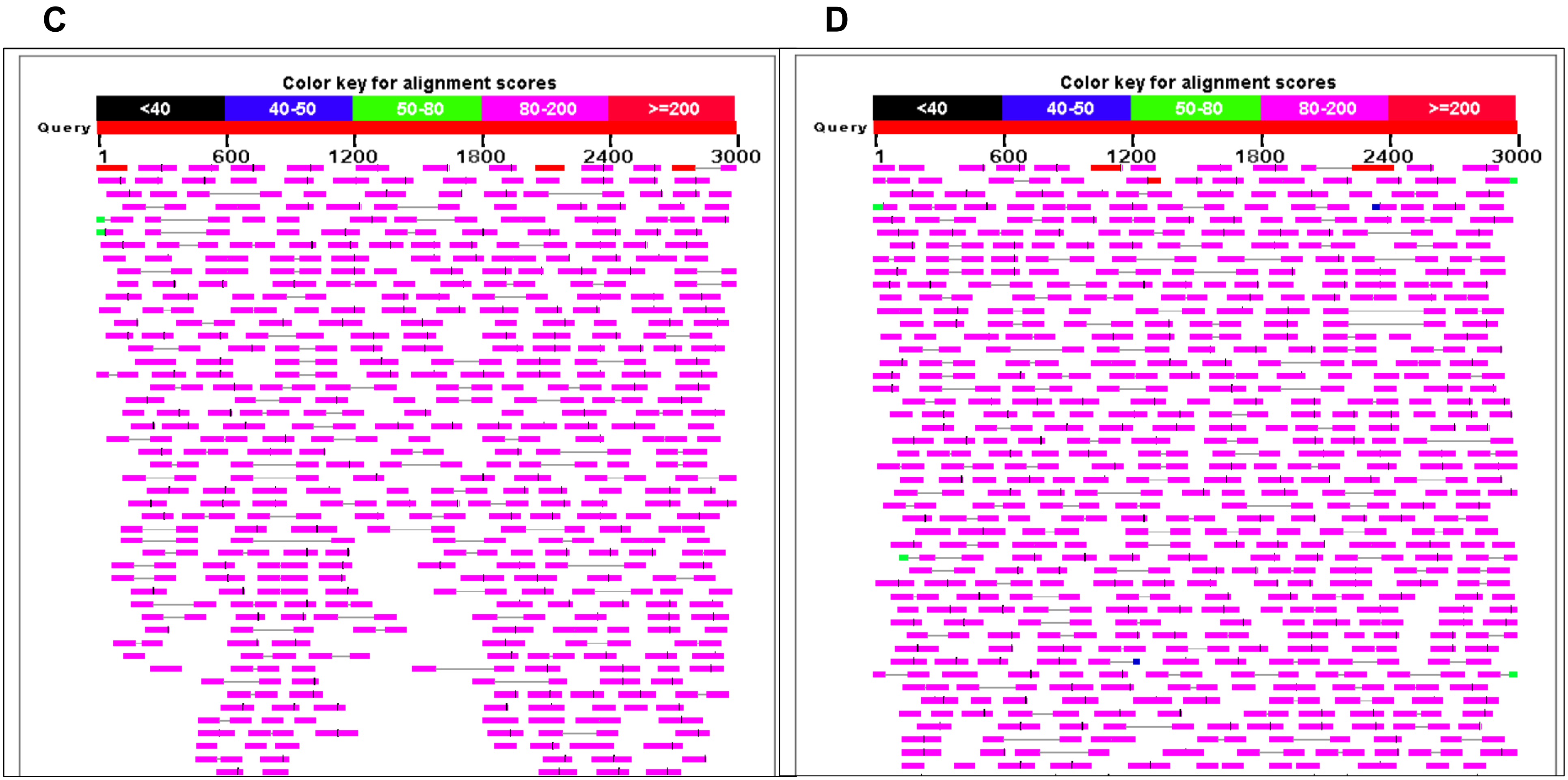


**Figure S5. RNA-Seq data for the "long" PMMs from *Lysobacter enzymogenes* and *Pseudomonas aeruginosa*.**
Graphical overview (1000 hits) of the outputs of MegaBLAST of 3-kb *dut-algC* DNA fragments from *Lysobacter enzymogenes* OH11 (Genbank: RCTY01000005, 9953..12952, panels A, B) and *Pseudomonas aeruginosa* UCBPP-PA14 (GenBank: CP000438, 6263694..6266694, panels C, D) against SRA entries SRX4941505 (A), SRX4941505 (B), SRX2549303 (C), and SRX2549304 (D). The *L. enzymogenes* PMM ORF starts at position 682, the *P. aeruginosa* ORF starts at position 436.
